# Intramolecular loops control SARS-CoV-2 nucleocapsid protein self-association and nucleic acid binding dependent on phosphorylation

**DOI:** 10.64898/2026.01.10.698783

**Authors:** Ai Nguyen, Siddhartha A.K. Datta, Camden Trent, Di Wu, Zillay Saleem, Ewa Szczesna, Wai-Ming Yau, Felicia Owoborode, Yan Li, Armin N. Adly, Heather R. Kalish, Grzegorz Piszczek, David O. Morgan, Peter Schuck, Huaying Zhao

## Abstract

The nucleocapsid protein of SARS-CoV-2 scaffolds genomic RNA into ribonucleoprotein complexes (RNP) for assembly in the virion, but also fulfills critical intracellular functions in replication and the suppression of host defense. It is comprised of a folded nucleic acid binding domain (NTD) and a dimerization domain, connected by a disordered linker containing a serine/arginine-rich (SR) region and a leucine-rich sequence (LRS). The switch between intracellular and assembly functions of N-protein is controlled by phosphorylation of the SR region, but the molecular details are unclear. Here we describe a model in which two mutually exclusive intramolecular loops bind the NTD and dynamically control self-association and nucleic acid binding properties dependent on the SR linker phosphorylation state. The model is supported by biophysical properties and interactions of full-length protein, point mutants, and peptide fragments. We find SR linker phosphorylation compacts the protein and inhibits nucleic acid binding and RNP formation, while enhancing self-association through promotion of transient coiled-coils in the LRS of the linker. These changes shift the nucleocapsid protein to a configuration poised for multi-valent interactions that support intracellular functions.

## INTRODUCTION

Intrinsically disordered regions (IDRs) in proteins support dynamic control and multifunctionality through a variety of mechanisms, including transiently folded motifs, functional switches controlled through post-translational modifications, fuzzy binding modes with multiple weak and spatially distributed intermolecular contacts, intramolecular loops, and liquid-liquid phase separation (LLPS).^1^ IDRs are particularly prevalent in RNA virus proteins,^2,3^ where they span a large biophysical parameter space across the quasispecies,^4–7^ support rapid evolution and efficient host adaptation through short linear motifs (SLiMs), and afford mutational tolerance in their interactions through fuzzy complexes.^8–10^ These features are exemplified by SARS-CoV-2 nucleocapsid (N) protein, which presents an excellent model system, not the least because the large mutational database of SARS-CoV-2 genomes provides a unique biophysical tool to elucidate protein functions and host adaptation mechanisms of an important viral pathogen.

N is the most abundant viral protein in the infected cell, constituting an estimated ≈1% of total protein.^11^ It fulfills a variety of functions related to host defense and viral replication, in addition to its critical structural role in the packaging of the ≈30kb RNA genome into ribonucleoprotein particles (RNPs) for viral assembly.^11–13^ N protein contains two folded domains – the N-terminal nucleic acid (NA) binding domain (NTD) and the C-terminal dimerization domain (CTD) (**Figure 1A**). ^14–18^ These are flanked and connected by long disordered regions (N-arm, linker, and C-arm) that cause much of the protein to be highly flexible and poised for dynamic regulation, for example, by allowing the central portion of the linker to make contacts across nearly the entire protein.^19^

**Figure 1:**
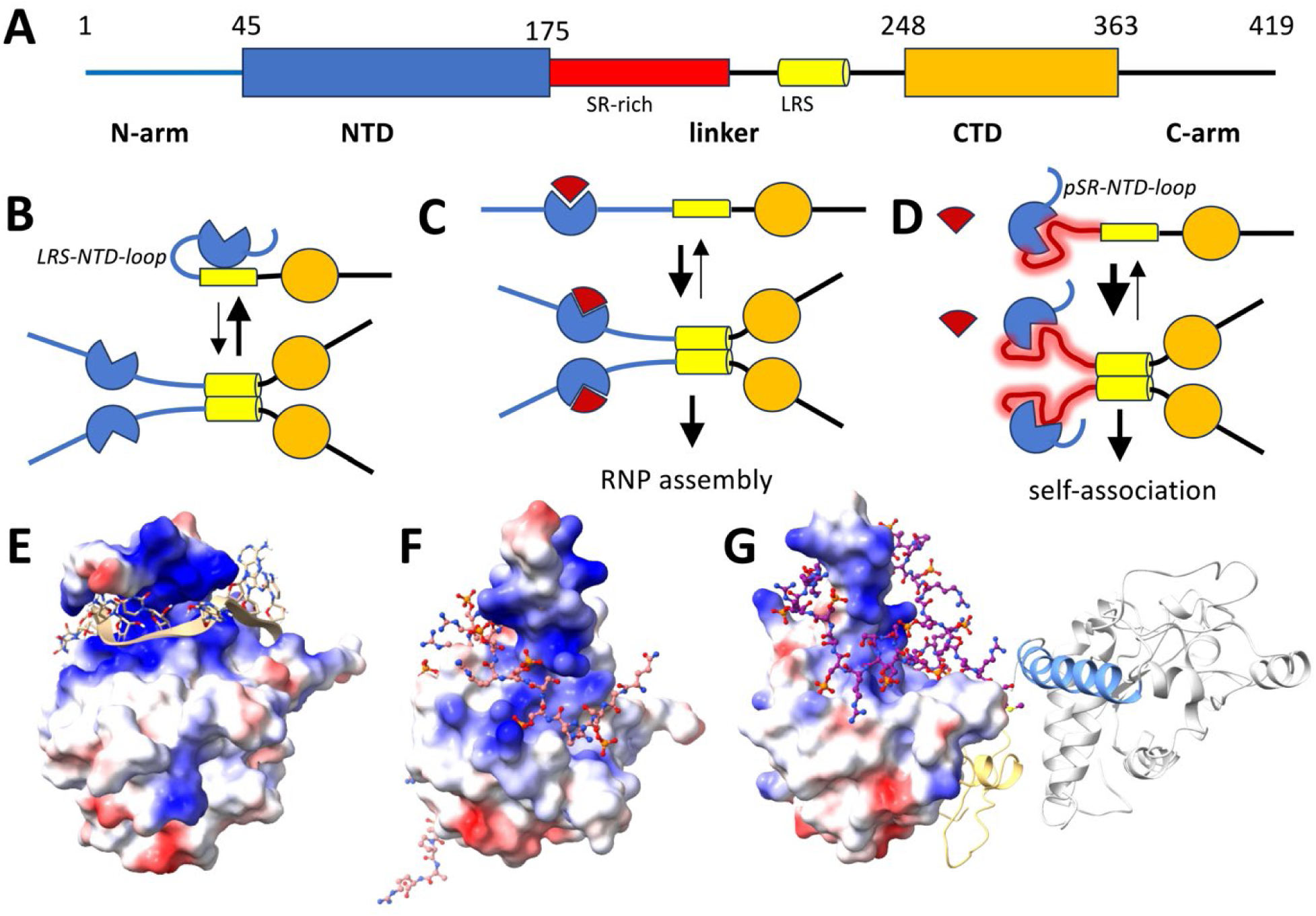
Schematic of N-protein organization and conformational states. (**A**) Subdivision of the 419 aa of N-protein in folded domains (NTD and CTD) and intrinsically disordered regions (N-arm, linker, and C-arm) that render the protein highly flexible, with a radius of gyration ranging from 4 to 8 nm.^19^ The linker (175-248) contains the SR-rich region (175-205, red) and a leucine-rich sequence that can form a transient helix (215-235, yellow) that allows promiscuous higher-order oligomerization. (**B**) Equilibrium of unmodified N-protein between dimeric and tetrameric states. For clarity only a single chain is drawn for each dimer, which would be linked with high affinity by the CTD.^16–18^ Tetramerization can occur transiently but is suppressed by the inhibitory LRS-NTD-loop, an intramolecular contact of the NTD (blue) with the LRS (yellow) and/or CTD (orange).^35^ (**C**) Binding of RNA (red sector) to the NTD releases the LRS and strongly promotes its assembly. NA-binding at multiple interfaces allows the formation of ribonucleoprotein particles (RNPs).^22,23^ (**D**) Phosphorylation of the SR-rich region (red shade) promotes binding of a loop to the RNA-binding site of the NTD (pSR-NTD-loop), competing with RNA binding and releasing LRS to self-associate.^35^ Formation of the pSR-NTD-loop and LRS-NTD-loop is mutually exclusive. (**E**) NMR structure 7ACT showing the NTD (shown with electrostatic surface) in complex with a 10mer ssRNA (ribbon and sticks).^14^ (**F**) AlphaFold3 (AF3) predicted structure of the NTD surface in complex with a phosphorylated peptide comprising the SR-rich region 181-210. (**G**) AF3 predicted structure of a contiguous pN including N-arm (yellow), NTD (electrostatic surface), phosphorylated SR-rich linker (sticks), LRS (blue) and CTD and C-arm (grey).

The C-terminal half of the disordered linker harbors a leucine-rich sequence (LRS) capable of transiently folding into helices that can weakly self-associate into promiscuous coiled-coils.^20,21^ Occupation of the NA binding site of the NTD induces a long-range conformational change that stabilizes the helical state of the LRS and enhances LRS self-association.^18,21^ Multivalent RNA is thought to bind multiple sites on NTD and CTD to help stabilize higher-order N oligomers (including hexamers of N dimers) into RNPs that scaffold the viral genome in the virions.^8,22–25^ While the allosteric mechanism that couples NA binding in the NTD to LRS self-association has remained elusive, NMR data corroborate the linkage, exhibiting broadened resonances in the LRS upon occupation of the NA binding site in the distant NTD.^26–28^

The central disordered linker also contains a serine/arginine (SR)-rich region adjacent to the NTD that displays 16 to 29 potential phosphorylation sites (dependent on mutant species^5^) and is strongly phosphorylated in cellular N-protein in infected cells ^29–31^ – in contrast to its state in the virion, where it is not or little phosphorylated.^31–33^ Phosphorylation occurs early during infection through a cascade of kinases and serves to inhibit NA binding and RNP assembly while promoting transcriptional and other cellular functions.^23,30,34–38^ These include the regulation of transcription of genomic and sub-genomic RNA,^32^ binding of 14-3-3 proteins^11,38–40^ and NSP3,^41^ and the modulation of N-protein condensation properties that facilitate partitioning of phosphorylated N-protein (pN) into stress granules.^29,34,42–46^ The phosphorylation-driven functional switch of N-protein functions has undergone evolutionary changes through defining mutations in most variants of concern, underscoring its importance in viral adaptation to the human host,^8,33,36,47–50^ and phosphorylation has been identified as a therapeutic target.^30,45,46,51,52^

The molecular basis for altered biophysical properties of pN is intensely studied but still incompletely understood.^23,35,36,44,45,53,54^ Recently, Botova *et al.* reported NMR data that reveal an intramolecular loop of the phosphorylated SR-rich linker region (pSR) occupying the NA-binding site of the NTD (a configuration we refer to here as pSR-NTD-loop), thereby competitively inhibiting NA binding to the NTD.^35^ Additional intramolecular contacts were found between the more distal linker region, as well as the CTD, and residues of the NTD outside the NA binding pocket (referred to as LRS-NTD-loop; see **Discussion**); these contacts were abrogated by phosphorylation.^35^ Previously, based on the observation of enhanced assembly of truncated N constructs lacking the NTD, Adly et al. hypothesized the existence of inhibitory intramolecular contacts localized at the NTD that control LRS oligomerization.^36^ Whether the LRS-NTD-loops identified by NMR constitute these inhibitory contacts, and how they couple phosphorylation with self-association and RNP assembly, remain unclear.

In the present work we synthesize these observations into a single mechanistic model of two distinct and mutually exclusive modes of intramolecular loops docking at the NTD, and thereby coupling phosphorylation or NA-binding to reversible self-association and higher-order assembly of N protein. The key features of the model are illustrated in **Figure 1B-C**. In the unphosphorylated unliganded state, the LRS-NTD-loop is partially occupied, thereby inhibiting the formation of coiled-coils in the LRS and N self-association (**Figure 1B**). If the unphosphorylated protein binds NA in the NTD, the LRS-NTD-loop is released allowing enhanced (unrestricted) LRS self-association (**Figure 1C**). Phosphorylation leads to the formation of the pSR-NTD-loop, which competes with NA binding to the NTD, but similarly releases the LRS-NTD-loop to enhance LRS self-association (**Figure 1D**).

To test this model, we examined the binding and assembly modes of N and pN, applying *in vitro* biophysical techniques including sedimentation velocity analytical ultracentrifugation (SV-AUC) to measure hydrodynamic shapes, macromolecular size-distributions, and protein-protein interactions. This is used to demonstrate direct binding of phosphorylated, but not unphosphorylated, SR peptide fragments to the NTD and full-length (FL) protein. Further, the presence of intramolecular loops is corroborated through measurements of thermodynamic stability of the NTD, hydrodynamic radii, and NA-binding of different phosphorylated or phosphomimetic protein constructs and mutants. While conflicting conclusions have been drawn in previous literature on the molecular properties of pN *vs.* N^35,45,53,54^, we will discuss how these can be largely resolved based on the new results^53^. The altered self-assembly properties associated with the different linker states contribute to the molecular basis for the switch between RNP assembly and cellular functions controlled by phosphorylation, and point to roles for N self-association in multi-molecular cellular complexes.

## RESULTS

### Phosphorylated SR-rich linker peptide binds the N-protein NTD in solution

The interaction between pSR and NTD previously discovered by Botova *et al.* in NMR chemical shifts of pN^35^ and predicted by AlphaFold3 (AF3) (**Figure 1G**) is the basis of our model in **Figure 1D**. We asked whether this interaction can be observed between dissected components free in solution. **Figure 2A** shows binding experiments by SV-AUC between the isolated NTD (N_48-173_) and a peptide fragment comprising the SR-rich region (N_175-208_). The isolated NTD sediments at *s*_w_ = 1.74 S (black), and a mixture of 40 μM NTD with a large excess of 100 μM unphosphorylated SR-peptide leads to virtually unchanged sedimentation of the NTD (*s*_w_ = 1.73 S; grey), identical within the precision of *s*-values of ≈0.01 S. By contrast, in mixture with enzymatically phosphorylated SR-peptide carrying 4 – 8 phosphate groups (pSR) a significant shift of the NTD sedimentation to higher values can be discerned (*s*_w_ = 1.85 S, red). This demonstrates direct binding of phosphorylated SR-peptide to the NTD free in solution, in contrast to the unphosphorylated SR-peptide, consistent with the AF3 prediction depicted in **Figure 1F**.

**Figure 2:**
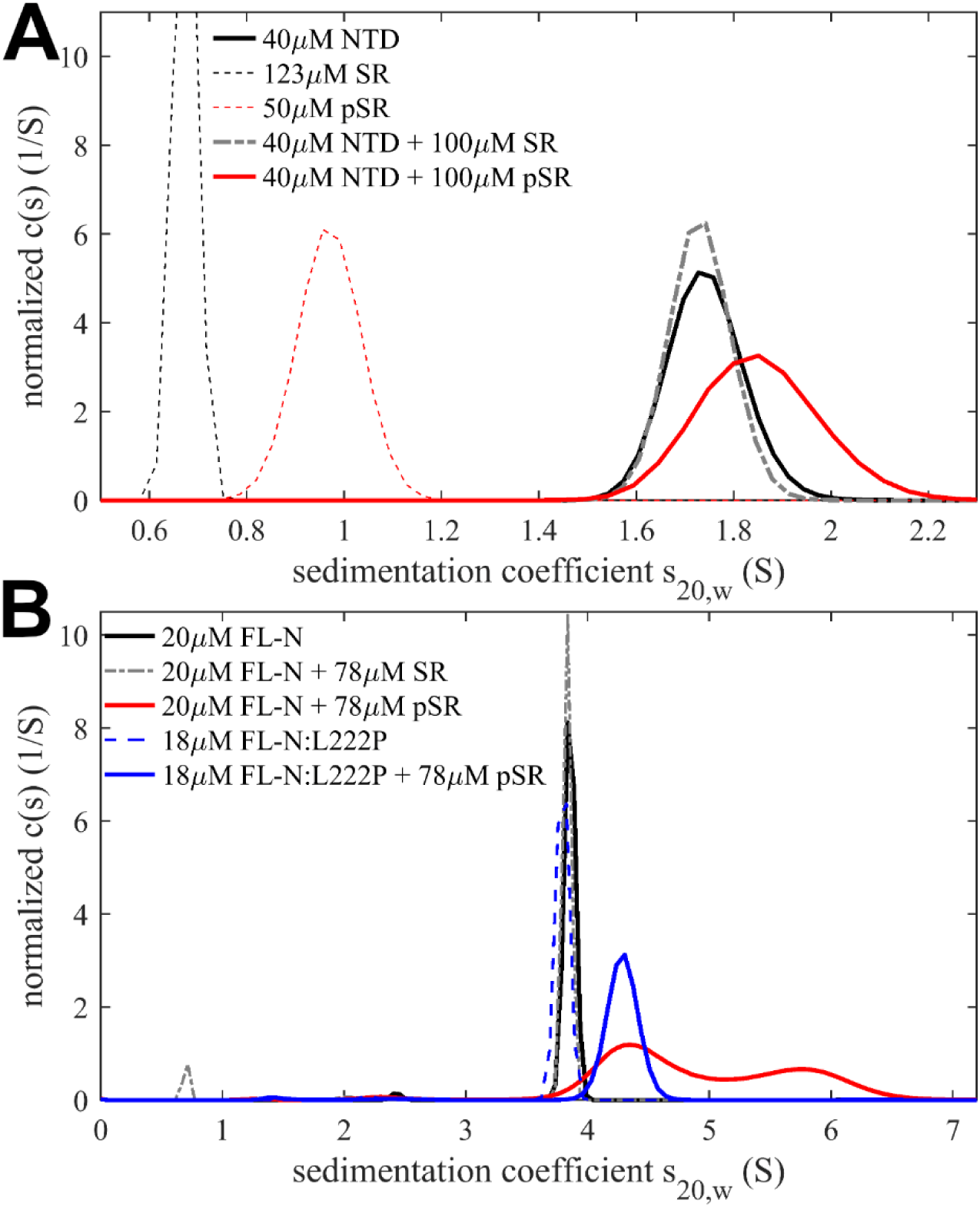
Binding of phosphorylated and unphosphorylated free SR peptide to NTD and full-length N in solution. SV-AUC experiments were carried out in 20 mM HEPES, 75 mM KCl, pH 7.50, acquiring sedimentation profiles by absorbance at 230 nm, monitoring largely the NTD and FL-N. Sedimentation coefficient distributions were normalized to unit area for comparison of individual components and peptide mixtures with isolated NTD (A) and full-length N (B).

A second element of our model is that occupation of the NTD binding site allows unhindered LRS oligomerization. This would imply that binding of pSR peptide to the FL N protein leads to higher-order self-association of N through the LRS. This was tested in the experiment of **Figure 2B**, which shows N does not interact with unphosphorylated SR, but sediments significantly faster in the presence of pSR. To dissect the role of LRS oligomerization from the effect of added mass of two bound peptides (total ≈8 kDa) on the sedimentation rate, we repeated the same experiment with the N:L222P LRS mutant that destabilizes the helical LRS state and abrogates LRS coiled-coil oligomerization. ^21^ In the presence of pSR, binding to N:L222P can be discerned from the *s*-value shift by ≈0.4 S (**Figure 2B**, blue), which is consistent with the added mass of pSR in the absence of oligomerization. By contrast, the observed shift of up to ≈2 S in the absence of the LRS mutation (**Figure 2B**, red) is consistent with the predicted N oligomerization and the formation of trimeric and/or tetrameric N species, similar to the previously reported size-distribution of N in the presence of short nucleic acid ligand T_10_ ^18,21^ .

The detailed energetics of free pSR peptide binding to NTD and FL N in solution was not pursued, as the interaction is likely significantly enhanced in the context of the FL protein where the SR-peptide ligand is already flexibly tethered to the NTD and complex formation thereby entropically favored. In the context of the FL N, the interaction of pSR with the NTD only requires a conformational change, for example, as predicted by AF3 in **Figure 1G**. For this reason, we turned to a detailed analysis of the phosphorylated and unphosphorylated FL N-protein.

### Phosphorylation of the SR-rich linker compacts N-protein

To study the effect of phosphorylation, we followed a previously described strategy in which bacterially expressed FL N-protein is incubated *in vitro* with the kinases SRPK1, GSK-3, and CK1, resulting in hyperphosphorylation of the SR-rich linker as previously characterized in detail by mass spectrometry mapping and NMR by several laboratories.^23,30,35,45^ We verified successful phosphorylation in our pN preparations by mass spectrometry, which showed the majority of molecules carrying 13-15 phosphate groups with a mode of 14 and only traces of other species (**Supplementary Figure S1A**). For comparison, we also expressed a phosphomimetic mutant FL N-protein in which 12 serine and threonine residues of the SR-rich linker region are replaced by aspartate (N_12D_), similar to the multi-D constructs of previous studies.^23,34,43^ However, it should be noted that each aspartate side chain provides only one negative charge at physiological pH, while each phosphate group carries two. Therefore, differences might be expected in the behavior of N_12D_ and pN (see **Supplementary Figure S1B**).^55,56^ All experiments in the present study were carried out at high nM to low µM N-protein, far above the dimerization dissociation constant *K_D,1-2_* of the CTD protein-protein interface, which is in the low nM range ^17,18^, such that the protein acts as a constitutive dimer. This likely is the functional unit in cells due to the high intracellular concentrations of N protein.^11^

Consistent with expectations for a largely disordered macromolecule, N-protein exhibits high translational friction.^18,57^ Therefore, the measurement of its sedimentation coefficient, which reports on the time-average of the conformational ensemble, can provide a sensitive measure for compaction or elongation. In the model of **Figure 1**, formation of the pSR-NTD-loop should lead to compaction, opposed by the possible extension from the release of the LRS-NTD-loop and the increase in size associated with weak LRS-mediated higher-order self-association (such as tetramerization); however, the latter is concentration-dependent while the former is not. **Figure 3A** shows isotherms of weight-average *s*-values as a function of protein concentration. Extrapolation to low concentration yields the *s*-value of the constitutive N dimer. This measurement is facilitated by the fact that experimental concentrations are low compared to the tetramerization dissociation constant *K_D,2-4_* (dimerization constant of two dimers forming a tetramer) while remaining far above the CTD dimerization constant *K_D,1-2_* (dimerization constant of two monomers to form a dimer). Furthermore, obligatory contributions from hydrodynamic nonideality that reduce the sedimentation velocity at high total macromolecular volume fractions in solution vanish in the limit of low concentrations.^58^

**Figure 3:**
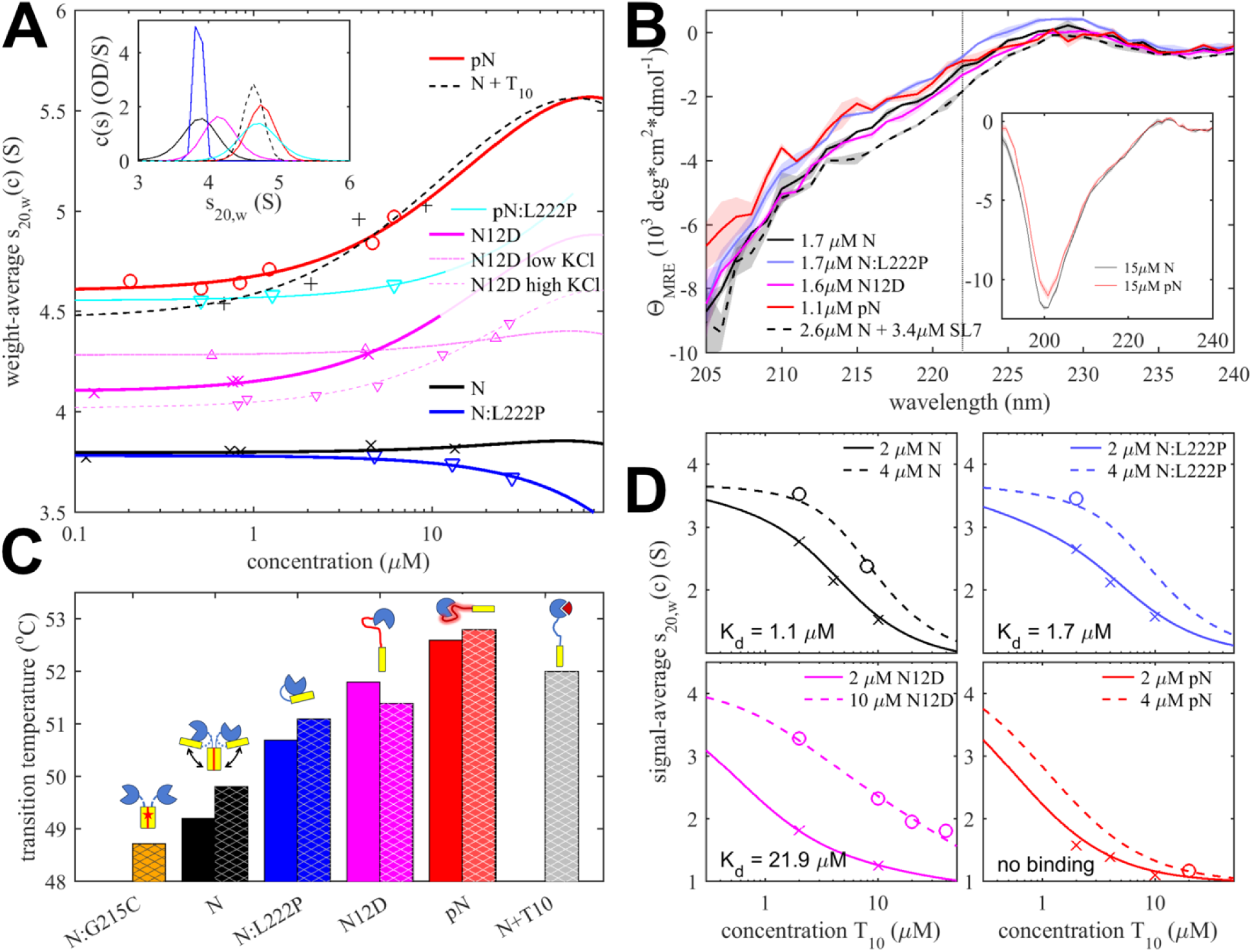
Impact of phosphorylation on structure, thermodynamic stability, self-association, and NA binding. (**A**) Compaction and enhanced self-association revealed by isotherms of weight-average sedimentation coefficients of different N constructs: N (black crosses), N:L222P (blue triangles), N_12D_ (magenta crosses), pN (red circles), pN:L222P (cyan triangles) and N in excess of T_10_ (black +) acquired in 20 mM HEPES, 75 mM KCl, pH 7.50. For comparison, N_12D_ isotherms are also shown in the same buffer with lower salt (10 mM KCl; magenta up triangles) and higher salt (150 mM KCl; magenta down triangles). Experimental precision is ≈0.01 S, and best-fit isotherms of a self-association model are shown as lines. The inset shows c(s) distributions at the lowest concentration in the corresponding isotherms. (**B**) Absence of increased helicity of pN and N_12D_, assessed by circular dichroism spectroscopy (CD) of N, N_12D_, pN, N:L222P, in comparison with the mixture of N with the stem loop 7 RNA molecule (SL7) that assembles into RNPs, in 20 mM HEPES pH 7.5, 75 mM KCl. The vertical line at 222 nm is indicative of helical content. To illustrate experimental error, the patches surrounding the spectra depict the wavelength-dependent standard deviation of 3 replicate scans. The inset shows spectra of N and pN extending further into the UV in low salt buffer (20 mM HEPES pH 7.5, 10 mM KCl). (**C**) Intramolecular loops are NTD ligands that enhance its thermodynamic stability. Transition temperatures observed by differential scanning fluorometry (DSF) reporting on the stability of the NTD for the different constructs at 3 µM, in 20 mM HEPES pH 7.5 with either 75 mM KCl (solid bars) or 150 mM KCl (hatched bars). Experimental precision is ≈0.2 °C. DSF scans are shown in **Supplementary Figure S4**. Cartoons are simplified depictions of the NTD ligation state, following Figure 1B-D. (**D**) Phosphorylation reduces affinity of the NTD for NA. Shown are isotherms of signal-weighted average sedimentation coefficients of mixtures of N, N:L222P, N_12D_, and pN with T_10_ in 20 mM HEPES, 150 mM KCl, pH 7.5 (symbols). Precision of the *s*_w_ data points is ≈0.01 S. Solid lines are best-fit isotherms with a model of two equivalent sites for T_10_ per N dimer.

The increase of the sedimentation coefficient from the unmodified value of 3.87 (3.85 – 3.90) S to 4.19 (4.16 – 4.21) S for N_12D_ and further to 4.70 (4.66 – 4.74) S for pN is highly significant (**Figure 3A** and **Table 1**). N_12D_ and pN have slightly higher masses (by 0.7% and 2.5%, respectively), but this cannot cause more than a proportional change in *s*-value, even in the conservative unphysical case that there was no accompanying opposing change due to increased molecular volume and buoyancy. Therefore, we can deduce that the observed >20% increase in *s*-value must be attributed to molecular compaction after phosphorylation, consistent with the significant population of the pSR-NTD-loop indicated by the NMR contacts observed by the Blackledge laboratory^35^, the prediction in **Figure 1G**, and the interaction between pSR and NTD observed in the direct binding assay of **Figure 2A**. Since the net effect is compaction, the release of the LRS-NTD-loop after phosphorylation either does not result in significant overall extension (i.e., the LRS-NTD-loop leads to a far less compact configuration than the pSR-NTD-loop), and/or the LRS-NTD-loop is significantly less populated in the unphosphorylated state under the explored experimental conditions.

**Table 1.** Compaction and interactions of phosphorylated and unmodified N-protein constructs.

| species | monomer mass (kDa) | dimer s-value<sup>(1)</sup> (S) | Stokes radius<sup>(2)</sup> (nm) | $K_{d,2-4}$ (μM dimer)<sup>(3)</sup> | $K_{d,NA}$<sup>(4)</sup> (μM) | DSF transition temperature 75 mM KCl ($^{\circ}$C) | DSF transition temperature 150 mM KCl ($^{\circ}$C) |
| --- | --- | --- | --- | --- | --- | --- | --- |
| N | 45.5 | 3.87 | 5.72 | 675 (401 – 1782) | 1.1<sup>(6)</sup> | 49.2 | 49.8 |
| N + T<sub>10</sub> | 45.5 + 3 | 4.56<sup>(5)</sup> | 5.17<sup>(5)</sup> | 36 (16 – 110) |  |  | 52.0 |
| N<sub>12D</sub> | 45.8 | 4.19 | 5.32 | 73 (66 – 107) | 21.9 | 51.8 | 51.4 |
| N<sub>12D</sub> + T<sub>10</sub> |  |  |  |  |  |  | 51.8 |
| N:L222P | 45.5 | 3.86 | 5.73 | >10,000 | 1.7 | 50.7 | 51.1 |
| N:L222P + T<sub>10</sub> | 45.5 + 3 | 4.56 | 5.17 |  |  |  | 52.8 |
| N<sub>12D</sub>:L222P | 45.5 |  |  |  |  | 53.0 |  |
| pN | 46.6<sup>(7)</sup> | 4.70 | 4.82 | 65 (53-84) | >250 | 52.6 | 52.8 |
| pN:L222P | 46.6 | 4.65 | 4.87 | 402 (186 – 542) |  |  |  |

<sup>(1)</sup> Limit of low concentrations in 20 mM HEPES, 75 mM KCl, pH 7.50 from isotherm analysis (**Figure 3A**), corrected to standard conditions. <sup>(2)</sup> Calculated based on extrapolated dimer s-value and mass. <sup>(3)</sup> Values in parenthesis are 68% confidence intervals.
<sup>(4)</sup> Binding of T<sub>10</sub> to the NTD measured by SV-AUC in high salt buffer (150 mM KCl) to suppress LRS oligomerization in the experimental concentration range. <sup>(5)</sup> Determined from N:L222P with saturating T<sub>10</sub>. <sup>(6)</sup> Previously reported.<sup>7</sup> <sup>(7)</sup> Mode measured by mass spectrometry.

It is possible to estimate the minimal number of residues that must be involved in the compaction. The hydrodynamic diameter of N is ≈11.4 nm, which will be reduced by 1.8 nm in the phosphorylated state. A fully stretched polypeptide would be shortened by this amount if at least ≈10 residues were folded over, assuming a peptide unit length of 0.38 nm.^59^ Considering that the SR-rich linker is unlikely to be fully stretched, we can conclude that a significant fraction of its ≈30 residues must change conformation in the pSR-NTD-loop to cause the observed hydrodynamic compaction.

Predictions by AF3 do not lead to a unique structure but consistently predict a substantial portion of the phosphorylated SR-rich linker to be folded into the NA binding groove of the NTD surrounding the base of the positively charged finger (**Figure 1F,G**) as seen in NMR structures of the NTD in complex with RNA^14^ (**Figure 1E**). This is consistent with the NMR chemical shifts of the NTD reported by Botova et al.^35^ upon N phosphorylation.

Interestingly, the hydrodynamic compaction of N_12D_ is only ≈40% of that of pN, which is suggestive of an incompletely formed, hydrodynamically less compact, or less populated loop. Since NA competition binding experiments lead to an estimated ≈95% occupancy of an NA-binding incompetent state of N_12D_ (see below), population differences cannot fully account for the lower *s*-value, and therefore structural differences between the pSR-NTD-loop in N_12D_ and pN are likely.

The variation in ionic strength leads to a variation of the limiting *s*-value of the N_12D_ (**Figure 3A**). The higher *s*-value in low salt and *s*-value closer to unmodified N in high salt is consistent with an electrostatic driving force to form the pSR-NTD-loop.

### Formation of the pSR-NTD-loop enhances LRS-based reversible N-protein self-association

As described above, weak oligomerization of the LRS helical state into promiscuous coiled-coils is a key protein-protein interface in the assembly of RNPs.^22,23^ It can be observed in SV-AUC, where the increase of the weight-average *s*-values with concentration is a sensitive measure of reversible self-association, slightly opposed by repulsive hydrodynamic nonideality. As previously reported, the N:L222P mutation disrupts the transient helix formation in the LRS region of the disordered linker and thereby abrogates LRS-mediated higher-order self-association of N dimers.^21^ Indeed, the *s_w_*-isotherm for this mutant shows a slight decrease of sedimentation velocity with concentration, which can be modeled well by accounting for hydrodynamic nonideality alone (**Figure 3A**, blue). By contrast, the unmodified wildtype N exhibits a near constant *s*-value, which reveals weak-self association compensating for the obligatory repulsive nonideality (**Figure 3A**, black). Quantitative analysis leads to a dimer-dimer dissociation constant *K_d,2-4_* = 0.68 (0.40 – 1.78) mM (**Table 1**).

Much stronger self-association of pN can be discerned from the increase in *s*-value at higher concentrations (**Figure 3A**, red). Quantitatively, the enhancement amounts to an order of magnitude improvement to *K_d,2-4_* = 65 (53 – 84) μM for pN, with N_12D_ assuming a similar value of *K_d,2-4_* = 73 (66 – 107) μM (**Figure 3A**, magenta). This is consistent with the release of the inhibitory LRS-NTD-loop caused by the occupation of the NA-binding pocket with the pSR-NTD-loop, thereby rendering the LRS free to oligomerize, as expected from the effects of free pSR peptide binding to N shown in **Figure 2B**.

As a control, phosphorylation of the LRS helix-abrogating mutant, pN:L222P, shows significantly weaker oligomerization (**Figure 3A**, cyan line), confirming the LRS to be the origin of the self-association of pN, similar to N. From the ≈10-fold higher *K_d,2-4_* of unphosphorylated N relative to pN, based on a simple two-state model of oligomerization-competent and -incompetent N (**Supplementary Methods**), we can estimate that for unphosphorylated N the LRS-NTD-loop is occupied ≈70% of the time.

To compare the enhanced self-association of pN to the previously reported allosteric enhancement of LRS-based self-association by NA binding to the NTD of unphosphorylated N, we use a model NA substrate the oligonucleotide T_10_. At low µM concentrations T_10_ binds virtually exclusively to the NTD and not the CTD (**Supplementary Figure S2**), and is of sufficient length to fill the NTD binding site but does not allowing scaffolding of multiple copies of N.^18^ Unphosphorylated N in the presence of an excess of an oligonucleotide T_10_ (**Figure 3A**, black dashed) self-associates with *K_d,2-4_* = 36 (16 – 110) μM, which is very similar to the *K_d,2-4_* value of pN, with possible small differences hinting at electrostatic repulsion in pN slightly opposing oligomerization.

We asked whether the release of the inhibitory LRS-NTD-loop in pN readily allows population of the helical state of the LRS, or if the LRS remains largely disordered in its free monomeric unassembled state. To address this, we carried out circular dichroism spectroscopy (CD) experiments at low μM protein concentrations, far below the *K_d_* for reversible LRS self-association measured above (**Figure 3B**). As we have shown previously, N protein spectra are dominated by a characteristic large negative peak indicative of disorder, but exhibit a distinct increase in the negative ellipticity at ≈222 nm upon assumption of the helical state in the LRS.^20,21^ While the CD spectra of pN and N_12D_ may be impacted by the different configuration of the SR-rich region, the negative ellipticity at ≈222 nm reporting on the helical content of pN and N_12D_ is similar to unmodified N and N:L222P (**Figure 3B**, solid lines) and does not show the increased helical signature observed for the assembly mixture of N in the presence of the stem loop 7 RNA (SL7) that, as previously reported, stabilizes the helical configuration in the RNPs (**Figure 3B**, dashed).^22^ This suggests that the ‘ground state’ of the LRS linker region in pN and N_12D_ remains disordered. Furthermore, CD experiments in low salt buffer transparent at lower wavelengths show less negative molar ellipticity at ≈200 nm for pN relative to N, suggesting reduced disorder for N with a phosphorylated SR-rich linker region, consistent with intramolecular binding.

Due to the electrostatic contributions from the charges in the SR-rich linker region, the oligomerization of N dimers would be expected to be salt-dependent. This was examined using the phosphomimetic N_12D_ (**Figure 3A**, magenta). Whereas under the standard conditions of 75 mM KCl used above (to be consistent with the *in vitro* RNP assembly assay) *K_d,2-4_* = 73 µM, in buffer with only 10 mM NaCl, we observe much weaker self-association with *K_d,2-4_* increasing to 1.1 mM (**Figure 3A**, dotted magenta line), suggesting enhanced electrostatic repulsion in the absence of charge screening. By comparison, the WT unphosphorylated N self-association at the same low salt conditions remains almost identical in 10 mM NaCl (*K_d,2-4_* = 0.76 mM^18^) and in 75 mM KCl (*K_d,2-4_ =* 0.68 mM). Thus, in low salt conditions the gain in dimerization affinity from the release of the inhibitory LRS-NTD-loop is smaller than the electrostatic repulsion introduced in the charged SR-rich region of N_12D_.

We also studied the effect of increasing the ionic strength to 150 mM KCl. In this case, K_d,2-4_ for N_12D_ was two-fold weaker than in 75 mM KCl, with *K_d,2-4_* = 148 µM (**Figure 3A**, dashed magenta line). This points to additional factors beyond simple screening of charges in the SR-rich region. In fact, as we have reported previously, the affinity of the protein-protein interface of the LRS helices that provides the driving force for the self-association is itself ionic strength dependent,^22^ likely due to stabilizing contributions of salt bridges of neighboring helices in the coiled-coil oligomers,^21^ leading to weaker LRS self-association in 150 mM NaCl compared to 10 mM NaCl. It appears this factor becomes limiting for the self-association of N_12D_ at high ionic strength.

Finally, to better compare our results with data from NMR laboratories studying N at pH 6.5, we measured the self-association of unmodified N in 50 mM sodium phosphate, 150 mM NaCl, pH 6.5 and obtained a best-fit estimate for *K_d,2-4_* of 0.26 mM (**Supplementary Figure S3**).

### Intramolecular loops increase NTD thermodynamic stability

To study the impact of the intramolecular interactions between the loops and NTD on the thermodynamic stability of the NTD, we pursued a measure of its melting point. The fact that aromatic amino acids of N are localized exclusively in the folded domains, notably including three tryptophan residues in the NTD, favors the use of differential scanning fluorometry (DSF). In this technique, the change in solvent exposure of aromatic amino acids upon temperature-induced unfolding reports on the local thermal stability of the folded domains. The CTD conveniently exhibits both higher stability and smaller intrinsic fluorescence changes upon melting than the NTD,^60^ such that the observed fluorescence changes at ≈50 °C can be attributed to the melting of the NTD. While DSF is a popular tool to detect extrinsic ligand binding, due to their binding energy usually stabilizing the protein structure, the same principles apply to intramolecular ligands. DSF traces of the different N species are shown in **Supplementary Figure S4** and their inflection temperatures T_i_ are depicted in **Figure 3C** and listed in **Table 1**.

As would be expected, addition of T_10_ at saturating concentrations to occupy the NA binding pocket on the NTD (**Figure 1E**) significantly increases the stability and raises T_i_ by 2.2 °C (**Figure 3C**, grey). On the other hand, introduction of the mutation L222P also leads to a significantly more thermostable NTD (**Figure 3C**, blue). L222P disrupts the LRS helix (which is significantly populated at the T_i_, as shown previously by temperature-dependent CD^18^) and thereby eliminates the possibility of LRS-LRS contacts competing with the LRS-NTD-loop contact to the NTD (**Figure 1B**). As a consequence, relative to the wildtype N-protein, N:L222P enhances the population of the LRS-NTD-loop, which leads to an increase of T_i_ of the NTD by 1.3 – 1.5 °C. (The combination of the L222P mutation and the presence of T_10_ stabilizes the NTD even further, with a T_i_ increase of 3.0 °C; **Table 1**.) Furthermore, as shown in detail previously, the G215C mutation of N that arose in the Delta variant of concern enhances the LRS coiled-coil oligomerization^20,21^ and lowers the transition temperature by ≈1 °C by reducing the occupancy of NTD by the LRS-NTD-loop (**Figure 3C**, yellow). ^7^

For pN we observe the stabilization of the NTD (relative to the unmodified N) by ≈3.2 °C (similar to an observation described by Gutmann et al.^37^), and by 2.6 °C for the phosphomimetic N_12D_ (**Figure 3C**, red and magenta, respectively). This change is consistent with the SR-rich linker serving as an intramolecular ligand by occupying the NA binding site on the NTD (**Figure 1D,F,G**).

For the phosphomimetic mutant, we observe slight further stabilization in the presence of T_10_ (**Table 1**), which is consistent with the competition of the pSR-NTD-loop and NA for the same site on the NTD (see below). Similarly, we also observe enhanced stability when abrogating the LRS oligomerization with N_12D_:L222P (**Table 1**), which would be expected to increase the occupation of the LRS-NTD-loop contact.

### Phosphorylation decreases nucleic binding affinity of the NTD through competitive population of the pSR-NTD-loop

Consistent with the pSR-NTD-loop mechanism binding into the NA binding site of the NTD, it was previously observed that phosphorylation reduces the NTD affinity for NA.^35,37^ This is recapitulated in the binding isotherms of **Figure 3D**, acquired under high salt conditions that suppress higher-order LRS-driven oligomerization in the concentration range probed. For unphosphorylated N this leads to an estimate of *K_d,T10_* = 1.11 (0.96 – 1.29) µM^7^ and similarly 1.7 (1.5-1.9) µM for N:L222P. By contrast, N_12D_ binds T_10_ ≈20-fold weaker with *K_d,T10_* = 20.9 (19.8 – 24.4) µM, and virtually no T_10_ binding was detected for pN (*K_d,T10_* > 246 µM; isotherm for no binding shown in **Figure 3D**; see **Table 1**).

If we consider the NTD to exhibit an equilibrium of two conformational states, one that is NA binding competent, and a second conformation where the pSR-NTD-loop blocks NA binding, then the ratio of the observed NA binding constants K_d2_/K_d1_ allows determination of the equilibrium constant between the two conformations, K_12_ = (K_d2_/K_d1_ - 1) (**Supplementary Methods**).^61^ In the case of the phosphomimetic mutant, the observed reduction of T_10_ binding implies 94.7% occlusion of its NTD binding site by the pSR-NTD-loop of N_12D_ corresponding to an energy ΔG_SR_ of –7.04 kJ/mol. In the case of fully phosphorylated pN, occupation by the pSR-NTD-loop is > 99.6% (with ΔG_SR_ < –13.2 kJ/mol). (It is interesting to consider these binding energies in the context of the free peptide binding experiments of **Figure 2**, which would imply K_d_-values < 5 mM for pSR – consistent with the observed binding data, and of ≈60 mM for a phosphomimetic version SR_12D_ – consistent with ultra-weak binding in SV-AUC shown in **Supplementary Figure S5**).

### Phosphorylated N-protein can form higher oligomers through LRS coiled-coils but not RNPs

The analysis of the isotherm of concentration-dependent weight-average sedimentation coefficients above showed enhanced self-association due to phosphorylation. A limitation of these measurements is that they reflect the time-averaged assembly state without resolving individual oligomeric species, which are obscured in SV-AUC due to their rapid reversibility^62,63^ and relatively low populations at practical protein concentrations. However, less quantitatively, the propensity to form higher oligomers up to the size of the RNPs in viral packaging can be visualized by mass photometry (MP) after chemical cross-linking (**Figure 4A,B**).^36^

**Figure 4:**
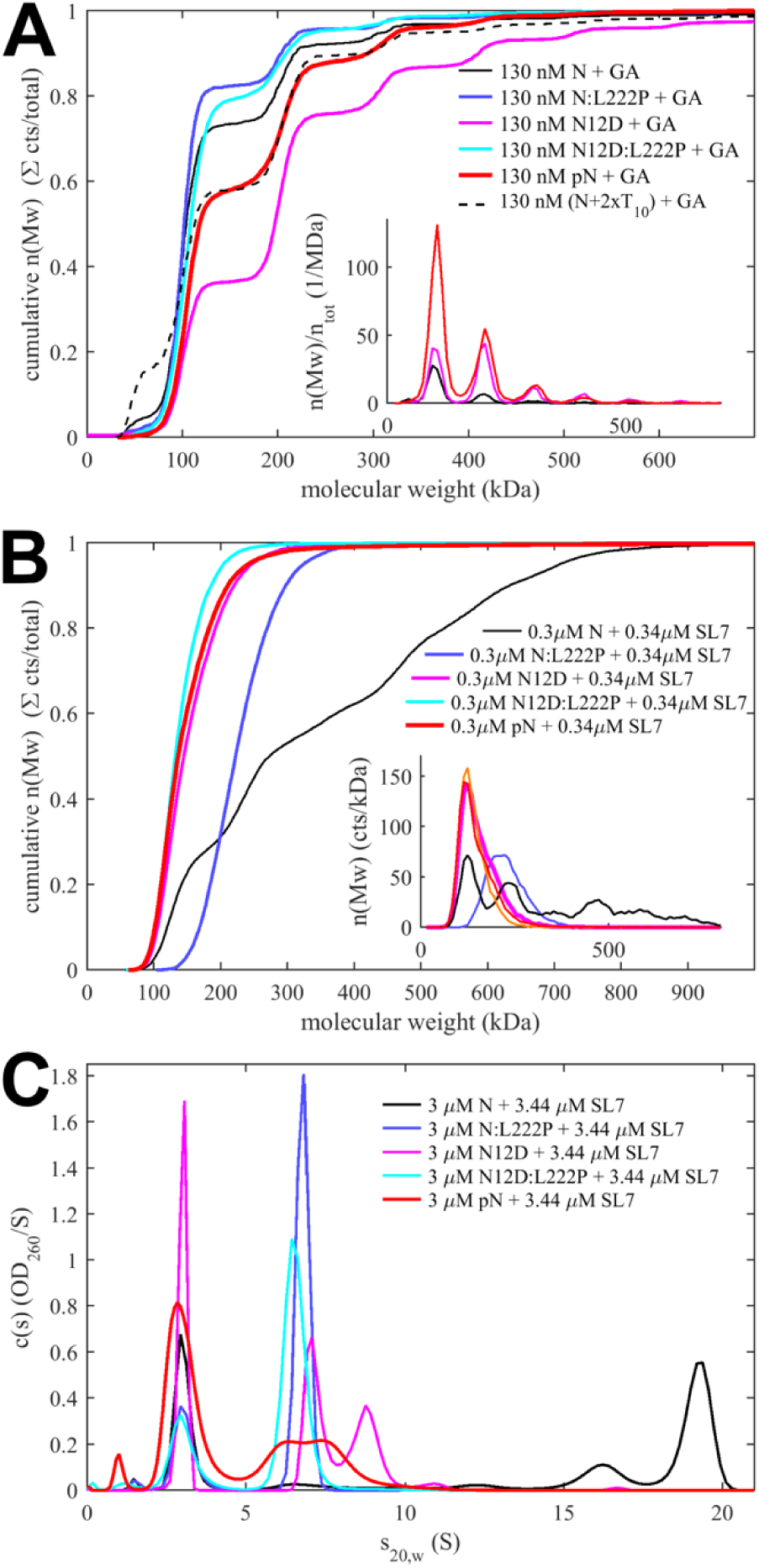
Impact of phosphorylation on higher-order oligomerization and RNP formation. (**A**) Propensity of higher-order oligomerization assessed by MP after crosslinking 15 µM N in 0.1% glutaraldehyde (GA) in 20 mM HEPES, pH 7.5, 75 mM KCl. Data are shown as cumulative molecular weight distributions, with selected normalized differential distributions as histograms in the inset. (**B**) In the absence of crosslinker, reversible formation of RNP particles of 0.3 µM N in assembly mixtures with 1.15-fold molar excess of SL7 RNA in 20 mM HEPES pH 7.5, 75 mM KCl. RNPs form in unmodified N but not in phosphorylated or phosphomimetic protein. (**C**) Sedimentation coefficient distributions measured at tenfold higher protein and RNA concentrations compared to the MP data shown in (**B**), showing abrogated RNP formation with modified N protein.

In the absence of NA ligands, unmodified N forms a ladder of higher oligomers including tetramers, hexamers, octamers, etc., in decreasing relative populations (**Figure 4A**, black). As a control, N:L222P forms virtually no oligomers beyond the dimer, confirming the LRS coiled-coil oligomerization as the assembly mechanism (blue). The polymerization ability is significantly enhanced in the phosphomimetic N_12D_ (magenta), in pN (red), and for N liganded with oligonucleotide T_10_ (black dashed), consistent with the improved availability of LRS coiled-coil self-association after displacement of the LRS-NTD-loop. That the abundant N_12D_ oligomers are indeed dependent on LRS self-association is demonstrated by their absence for the N_12D_:L222P mutant that abrogates LRS helices (cyan). This result shows that not only tetramerization, but higher-order N-protein self-association through the coiled-coil interaction of LRS helices is enhanced relative to unmodified protein due to phosphorylation.

As we have shown previously, this mechanism is key for the assembly of RNPs, where higher oligomers are further stabilized by additional ultra-weak protein-protein interfaces and multi-valent protein-NA interactions with suitable dsRNA substrates, such as stem loops from the 5’-UTR.^8,22,23^ Without chemical cross-linking, the addition of stem loop 7 (SL7) to N leads to the formation of 600 – 700 kDa RNP particles that may serve as models for RNP^8,22,23^ (**Figure 4B**).

Strikingly, the formation of such RNPs is completely abolished under the identical conditions for both the phosphomimetic N_12D_ and the enzymatically phosphorylated pN, limiting complexes to at most a tetramer at the sub-µM concentrations required in MP (**Figure 4B**). This is remarkable since unavailability of NTD as a NA binding site alone does not prohibit formation of RNPs, as published experiments with NTD deletion mutants (N*) show (see **Discussion**).^36^ Interestingly, even the tetramers of pN and N_12D_ are formed with SL7 at lower populations compared to the N:L222P mutant with abrogated LRS self-association (**Figure 4B**, blue), which suggests that at low concentration scaffolding of N on multiple NA sites is the main mechanism of tetramerization, which is weakened by phosphorylation. At tenfold higher concentrations in SV-AUC (**Figure 4C**), tetramer and higher oligomers with ≈7 – 10 S are formed by N_12D_ and pN in the presence of SL7, exceeding the tetramerization of N:L222P from scaffolding alone, consistent with concentration-dependent LRS oligomerization. As in the cross-linking assay of **Figure 4A**, the oligomers for N_12D_ are larger than those of pN, but far below the size of RNPs of unmodified N.

Another assay to assess weak self-association is *in vitro* condensate formation. Multi-valent protein-protein interactions generated by LRS coiled-coils and/or beta-sheet interactions in N-arms of the Omicron P13L mutants, ^8,21,43^ in conjunction with NA interactions, promote the formation of condensates with material properties dependent on the N phosphorylation state. ^34,43–45,53^ In the presence of a short oligonucleotide T_40_ that can scaffold only a few copies of NTD and CTD,^22^ we observe strongly reduced droplet formation for the phosphorylated and phosphomimetic variants pN and N_12D_ (**Supplementary Figure S6**). We attribute this to the reduction of NA interactions, highlighting their role in condensate formation of N.

## DISCUSSION

Phosphorylation of IDRs is a common motif regulating function through altered conformational ensembles that can modify transient folding, intramolecular loops, oligomerization, LLPS, and protein or NA binding.^56,64,65^ In the case of SARS-CoV-2 N-protein, it has long been recognized that phosphorylation of the SR-rich region in the central disordered linker constitutes a critical functional switch between assembly functions and intracellular functions, including viral replication and remodeling of host cellular processes, but the molecular mechanistic details have remained obscure.^31,34,44^

Several recent observations have pointed to the existence of phosphorylation-controlled intramolecular loops of the central disordered linker contacting the NTD, including long-range allostery between NA binding in the NTD and conformational changes and oligomerization of the distal linker LRS,^18,26,27^ altered RNP assembly functions of different N constructs,^36^ and most recently, NMR data from the Blackledge laboratory^35^ that reveal the phosphorylated SR-rich linker region is binding into the NA binding pocket of the NTD (the pSR-NTD-loop), mutually exclusive to additional, more long-range loops of the linker further downstream, and of the CTD, contacting other regions of the NTD (the LRS-NTD-loop). The pSR-NTD-loop provides a possible explanation for additional observations reported in the literature, including observed protection of N from limited trypsin digestion after phosphorylation,^54^ as well as the protection by the NTD against phosphatase activity of NSP3 dephosphorylating the linker.^66^ In support, previous molecular simulations of the N-protein conformational ensemble by Różycki & Boura have shown that the linker is extremely flexible and that the LRS can make frequent contacts with the NTD.^19^ Recent studies by Stuwe et al.^53^, Sullivan et al. ^54^, and Favetta et al.^45^ have contributed mechanistic aspects of the phosphorylation switch but not yet considered these loops.

In the present work, we describe new evidence for the direct interaction between pSR and the NTD, systematically examined further structural, hydrodynamic, and thermodynamic evidence of the pSR- and LRS-NTD-loops and their switch through phosphorylation, and explored functional consequences regarding self-association, NA binding, RNP formation, and LLPS.

First, we show that the interaction between phosphorylated SR-rich region is sufficiently strong that even an untethered, soluble pSR peptide fragment binds the NTD, and induces N oligomerization similar to nucleic acid ligands (**Figure 2**). In the context of the contiguous full-length N-protein this interaction results in an intramolecular loop. Our data show that phosphorylation leads to a ≈20% reduction of the N-protein Stokes radius, and thermodynamically stabilizes the NTD similar to the binding of NA, which it strongly abrogates (**Figure 3**). This is consistent with the formation of the pSR-NTD-loop where the phosphorylated SR-rich region binds into the NA binding pocket of the NTD, as indicated by chemical shifts in the NTD and pSR reported by Botova et al.^35^ An estimate of the number of residues involved in the loop, based on the change in overall molecular dimension, suggests a significant fraction or even the majority of residues in the SR-rich linker may be engaged in the loop, as suggested by predicted AF3 structures (**Figure 1F,G**).

Even the phosphomimetic N_12D_ variant shows ≈95% occupancy of the NA-binding pocket, although enzymatically phosphorylated pN binds significantly stronger, as judged by the virtually complete competition of NA binding by pSR. In this context, it is noteworthy that the fractional change in Stokes radius is much smaller for N_12D_ than that of enzymatically phosphorylated pN, which suggests that the NA-binding competitive state of the phosphomimetic mutant is different from that of enzymatically phosphorylated pN – likely a consequence of the lower charge density and/or other previously described imperfections of the phosphomimetic model.^55,56,67^ Nonetheless, the formation of the pSR-NTD-loop is strongly supported even with the N_12D_ construct, by still significant observed compaction and by the significant increase of the thermodynamic stability of the NTD (for N_12D_ exhibiting already 2/3 of the shift in T_i_ of pN), as would be expected for an intra-molecular ligand binding event.

The detailed pattern and sequence of phosphorylation have garnered significant attention.^30,35,54^ Botova et al.^35^ carried out detailed analyses of the kinetics of phosphorylation events by NMR. They observed no inhibition of RNA binding – and presumably no pSR-NTD-loop formation – after phosphorylation with PKA or SRPK1 alone, whereas SRPK1/GSK-3 completely abrogates RNA binding. Even with phosphomimetic mutants, Sullivan et al. ^54^ reported differences in the effect of C-terminal or N-terminal placement of three aspartate substitutions in their phosphomimetic constructs.^54^ This appears to suggest that phosphorylation of specific residues may be required for pSR-NTD-loop formation – a possibility supported by a comparison of consensus sequences of related coronaviruses where most linker serine and arginine residues coincide.^20,30^ However, a more detailed sequence analysis of the mutant spectrum of SARS-CoV-2 genomes in the GISAID database reveals that the SR-rich region is extremely variable:^20^ With the exception of S176, which is the sole conserved serine in the mutant spectrum, no specific serine or arginine residue is essential for infectious virus, and probably for the phosphorylation mechanism of N.^20^ (Notably, this includes S188 and S206, the priming sites by SRPK1.) Nonetheless, all sequenced SARS-CoV-2 species do maintain a minimum number of 6 SLiMs for GSK-3, 5 SLiMs for CK1, and 2 SLIMS for PKA at various locations in the linker.^5^ This suggest that a threshold local charge density may be essential. An in-depth analysis of the pattern of phosphorylation sites within the mutant spectrum may reveal additional specific structural or sequence requirements. If a threshold charge density is critical for the phosphorylation switch and formation of the pSR-NTD-loop, and, as it appears, the enzymatically phosphorylated N-protein with 13-15 phosphate groups is incompletely mimicked by the N_12D_ mutant, then such a threshold may not be reached for the much less charged phosphomimetic ‘6D’ and ‘3D’ mutants studied by Sullivan et al.^54^ In conjunction with methodological differences, this may explain less significant changes in self-association and thermodynamic stability observed in the Sullivan et al. study. ^54^

A second key finding of our study is the enhancement of higher-order self-association of N-protein upon phosphorylation, corroborating the release of the inhibitory LRS-NTD-loop in our model once the pSR-NTD-loop is formed (**Figure 1**). Clearly the LRS-NTD-loop is a simplified picture, since two long-range loops were indicated by the NMR data of Botova et al.^35^ However, we may consider all intramolecular contacts of residues downstream from the SR-rich region with the NTD to be different sub-states of the “LRS-NTD-loop”, to the extent that they inhibit LRS oligomerization and release upon occupation of the NA binding groove of the NTD through either phosphorylation or NA binding. This operational definition of the LRS-NTD-loop is sufficient for the functional and energetic characterization of the linkage between phosphorylation, NA binding, and LRS self-association even if it reflects an ensemble of looped microstates. Further studies are required to resolve higher molecular detail, including a more precise delineation of the binding interface on the NTD. Similar configurational ensembles may exist for the pSR-NTD-loop.

The enhanced LRS self-association observed in the present work is ostensibly inconsistent with the report by Stuwe et al.,^53^ who concluded from SV-AUC and NMR data that self-association of the LRS is reduced by phosphorylation, not enhanced. However, the discrepancy disappears considering that Stuwe et al. used constructs lacking the NTD,^53^ thereby missing both loops, specifically the LRS-NTD-loop that reduces baseline self-association in the unphosphorylated state. In addition, it appears that the lack of the NTD may permit non-native intra- or inter-molecular contacts of pSR, for example, with the NA-binding site of the CTD (as depicted in an AF3 prediction of **Supplementary Figure S7**). Indeed, SV-AUC experiments reveal binding of a phosphorylated pSR peptide to an isolated CTD construct in solution (**Supplementary Figure S8**). The latter is consistent with the observation by Adly et al.^36^ that RNA binding to an N construct lacking the NTD is inhibited by pSR. The lack of the NTD jointly with possible non-native intramolecular interactions in the resulting construct render the results by Stuwe et al.^53^ difficult to compare with results presented here.

Previous studies have compared the radii of gyration of phosphorylated and unphosphorylated N using SAXS, with conflicting results.^35,45^ A methodological limitation in this technique is the distinction between compaction and the enhanced self-association we describe in the present work. For example, the apparent lack of compaction in the SAXS experiments by Botova et al.^35^ might be related to the enhanced self-association of pN, which would tend to increase the radius of gyration and mask compaction. By using concentration series in SV-AUC in our study, exploiting its greater dynamic range, self-association can be quantified with high sensitivity and discriminated from hydrodynamic compaction. Furthermore, use of the L222P LRS-mutant allows selective inhibition of self-association. We also observed that the magnitude of self-association depends not only on concentration, but also strongly on buffer conditions. This might explain why Favetta et al. (in contrast to Botova et al.^35^) did observe significant compaction by SAXS,^45^ consistent with our hydrodynamic results. However, in conjunction with smFRET data of the linker and coarse-grained simulations, this was interpreted to arise not from pSR-NTD-loops to the NTD, but from new intra- or inter-molecular contacts.^45^ At least in part, this seems to be consistent with our finding of enhancement of LRS self-association upon phosphorylation which in our model arises through formation of pSR-NTD-loop and simultaneous release of the of the LRS-NTD-loop. Again, such altered weak reversible LRS-based self-association can be conclusively demonstrated by SV-AUC through the experiments with N:L222P LRS-abrogating mutants, but would have remained undetectable in the study by Favetta et al.^45^ Unfortunately, the strong electrostatic contributions to the various binding interfaces limit the ability to quantitatively compare experimental results at different pH and salt concentrations.

Once the LRS-NTD-loop is released, it appears the LRS still remains largely in an unfolded configuration, only transiently populating a minority helical state that is the prerequisite for oligomerization. We draw this conclusion based on the similarity of the circular dichroism spectra of N, N_12D_, and pN in comparison to spectra with significantly higher helical content under conditions where the LRS helices populate and provide self-association interfaces, such as in the N:G215C mutant from the Delta variant of concern^7,20,21^, or assembled in the RNP^22^. Intra-dimer LRS helix interactions appear to be rare events, consistent with molecular simulations of the N-protein conformational ensemble by Różycki & Boura that show low intra-dimer contact frequencies of the LRS with each other compared to contacts of LRS with the folded domains.^19^ It is possible that after release of the LRS-NTD-loop, cooperative assembly of LRS oligomers larger than the dimer is required for stabilization, which may occur in N-protein tetramerization and is likely a key contribution to the assembly of RNPs.^22^ In any case, based on the similarity of measured binding constants *K_D,2-4_*, one would expect that at this point the self-assembly process should be largely independent of whether the LRS-NTD-loop was previously released due to NA-binding or pSR-binding to the NTD.

On the other hand, if the only effect of the pSR-NTD-loop were to functionally eliminate the NTD as a NA binding site for stabilization of RNPs, and to free the LRS from its inhibitory interaction with the NTD, then it might be compared to a construct N* lacking the N-arm, NTD, and SR-rich region (N_210-419_, which arises in virus-infected cells as an alternate expression product from the N:R203K/G204R mutation^47,48,68,69^). Remarkably, however, N* has been shown to be capable of assembling RNPs,^36^ which are not observed for pN or N_12D_. (As described in ^23^ non-compact RNP-sized complexes of pN can be isolated only after stabilization through glutaraldehyde crosslinking.) We speculate that other factors that do not apply to N* contribute to the inability of pN to form stable RNPs, including charge repulsion in higher oligomers ^36,53^ (which would explain their larger populations for N_12D_ relative to pN in **Figure 4A** and **C**), electrostatic interactions of pSR affecting RNA binding to the CTD, and/or steric restrictions of the compacted full-length N configuration inhibiting assembly in the same mode as N*.

The fact that LRS interactions are enhanced and thereby N-protein self-association is strengthened after phosphorylation raises questions about the role of N-protein self-association in its intracellular functions. Even though pN blocks the NTD binding site for NA, which has a preference for single-stranded NA and is capable of binding either DNA or RNA, it still carries two NA binding sites in the CTD that have a preference for dsRNA.^22,70–73^ While the reduced multi-valency of NA binding critically diminishes the stability of RNPs, it is conceivable that the enhanced self-association of pN facilitates the dynamic assembly of multi-protein complexes involving multiple copies of pN. We speculate that this ability might modulate interactions with NSP3, chaperone activity, and transcription functions.

## METHODS

### Protein expression and phosphorylation

Full-length wild-type and mutant SARS-CoV-2 N-proteins were expressed and purified as previously described.^7,22^ Briefly, a pET29a(+) plasmid—containing a kanamycin-resistance gene and the gene encoding the N-protein of interest with an N-terminal 6xHis tag and a Tobacco Etch Virus (TEV) cleavage site—was transformed into One Shot BL21(DE3)pLysS E. coli (Thermo Fisher Scientific, Carlsbad, CA). The expressed protein was purified using Ni^2+^ affinity chromatography and underwent on-column unfolding and refolding to remove protein-bound cellular NA (which promotes artificial oligomerization and condensation).^34,74,75^ Following TEV protease cleavage, tag removal was verified through a second round of affinity chromatography and/or mass spectrometry before final size-exclusion chromatography. The purified protein was dialyzed into a working buffer (20 mM HEPES, 75 mM NaCl, pH 7.50). Protein purity was confirmed via SDS-PAGE, and the absence of NA was verified by an A260/A280 absorbance ratio of approximately 0.50–0.55. Final protein concentrations were determined by UV-Vis spectrophotometry or refractive index-detected SV-AUC. Protein homogeneity in size, shape, and oligomeric state was further confirmed by the sedimentation as a single *c*(*s*) peak in SV-AUC (e.g., **Figure 2B**). A construct comprising the NTD (N_48-173_) was similarly bacterially expressed and purified by Ni-NTA chromatography, as described previously ^22^.

N protein and mutants were phosphorylated *in vitro* as described.^36^ Protein kinases were acquired from Promega (SRPK1: #VA7558, GSK-3β: #V1991, CK1ε: V4160). Briefly, 16.5 μM N protein was mixed with 80 nM SRPK, GSK3, and CK1 in kinase reaction buffer (20 mM HEPES pH 7.5, 70 mM KCl, 10 mM MgCl_2_, 1 mM DTT, 0.5 mM ATP). The reactions were incubated at 30°C for 30 min. Alternatively, 41 μM N protein was mixed with 80 nM SRPK, GSK3, and CK1 in kinase reaction buffer (20 mM HEPES pH 7.5, 70 mM KCl, 10 mM MgCl_2_, 1 mM DTT, 1 mM ATP), followed by incubation at 37°C for 20 hrs. The reactions were quenched with 5 mM EDTA, and followed by dialysis into working buffer and size-exclusion chromatography. The extent of phosphorylation was monitored by mass spectrometry.

For the peptide consisting of the SR (N_175-208_), priming of serines 188 and 206 was carried out by synthesis with phosphoserine (ABI Scientific, Sterling, VA), and enzymatic phosphorylation was carried out by GSK3 and CK1 only, leading to 4-8 phosphate groups per peptide. The reaction mixture was purified of enzymes and free ATP by size-exclusion chromatography (Superdex 30, Cytiva, Marlborough, MA) followed by concentration through centrifugal filtration (Pierce MWCO 3kDa, Thermo Fisher Scientific).

### Sedimentation velocity analytical ultracentrifugation

SV-AUC experiments were performed in a calibrated ^76^ ProteomeLab Xl-I analytical ultracentrifuge (Beckman Coulter, Indianapolis, IN) following standard protocols ^77^. Briefly, AUC cell assemblies comprising 12- or 3-mm charcoal-filled Epon double-sector centerpieces were filled with the working buffer and indicated samples in the reference and sample sectors respectively. The AUC cell assemblies were then inserted into An-50 or An-60 rotors, followed by temperature equilibration at 20 °C for 2-3 hrs. Radial scans were collected at 50 krpm with Rayleigh interference optics and absorbance optics at 230 nm, 260 nm and/or 280 nm. Analyses of SV-AUC data were performed in the software SEDFIT using the sedimentation coefficient distribution model *c*(*s*).^78,79^ Distributions were integrated in GUSSI^80^ to determine signal-weighted average *s*-values, and isotherms were globally fitted in SEDPHAT using a self-association or ligand-linked self-association model, respectively. The N tetramer *s*-value was linked to that of the dimer by the hydrodynamic scale relationship *s*∝*M^2/3^*, ^62^ and nonideality was measured for N:L222P in the absence of self-association and in first approximation taken to be equal for all constructs.

### Mass photometry

MP experiments were carried out on a TwoMP instrument (Refeyn, UK) as previously described.^22^ The samples were prepared by diluting the stock solutions with the working buffer prior to MP data acquisition. For the MP experiment with crosslinked material (**Figure 4A**), N-protein was incubated at 15 µM protein with 0.1% glutaraldehyde for 10 min.^36^ A 9 µL volume of working buffer was added onto the microscope coverslip which was loaded on the microscope, followed by focusing, addition of 1 µL of sample to the buffer droplet, and gentle mixing. MP data was collected immediately afterwards using the AcquireMP software, and the analysis was performed with the DiscoverMP software (Refeyn, U.K.). The measured interferometric contrast values were converted to molecular weights by following the protocol provided by the manufacturer using β-amylase and thyroglobulin standards (Sigma Aldrich, St. Louis, MS, A8781 and T9145).

### Circular dichroism

CD spectra were collected in a Chirascan Q100 instrument (Applied Photophysics, UK) at 20 °C for the samples in the indicated buffers. Measurements were carried out in 1 mm or 0.1 mm pathlength cuvettes with 1 nm steps, and a 1 sec integration time per data point. For some measurements, adaptive sampling mode was used, in which the time per data point is varied inversely as a function of the detected light. For each CD spectrum, three independent scans were averaged following background subtraction. The CD signals were normalized to protein concentration, which was calculated from the absorbance at 200 nm or 205 nm of each sample, using a reference protein sample, whose concentration was determined by absorbance at 280nm with a UV-Vis spectrophotometer.

### Differential scanning fluorometry

DSF measurements were carried out in a Tycho instrument (Nanotemper, Germany) as previously described.^7^ Each sample (≈10 µL) was loaded into a capillary (TY-C001, Nanotemper, Germany), and fluorescence signals at 350 nm and 330 nm generated from the intrinsic fluorescence of the proteins were measured during a temperature ramp from 35°C to 95°C at a rate of 30°C/min. The inflection temperature (*T_i_*) of each sample is calculated in the Tycho software, as a measure of the transition temperature of the protein’s thermal unfolding process.

### Optical microscopy

Brightfield light microscopy of *in vitro* LLPS was performed as previously described.^7^ Images were acquired using a Nikon Ts2R inverted microscope equipped with a CF160 Plan Fluor 100X NA 1.3 oil objective lens and recorded with a photometrics moment camera with a pixel size of 44 nm. Samples were prepared by mixing 10 µM N protein and 5 µM T_40_ in a low salt buffer (20 mM HEPES, 10 mM KCl, pH 7.50), followed by imaging at room temperature.

### Structure prediction

Structure predictions were carried with AlphaFold on the AF3 server^81^. For the complex of the NTD (N_44-180_) with the phosphorylated SR-rich region 181-210 (pSR), all serines were specified as phosphorylated and the pSR and NTD were considered separate chains. For the prediction of the contiguous pN, four chains of pN (with all serines in the SR-rich region specified as phosphorylated) were assumed. This led to the prediction of a pN tetramer of which for clarity only a single pN chain is shown in **Figure 1G**.

### Mass spectrometry

The number of phosphate groups attached to N protein was measured by LC/MS or MALDI-TOF mass spectrometry.

LC/MS was carried out on an Agilent AdvanceBio 6545XT LC/Q-TOF Mass Spectrometer (Agilent Technologies, Santa Clara, CA) coupled with Infinity II HPLC. 100-200 ng of each protein in 10% acetonitrile/0.1% formic acid in water were injected to PLRP-S 1000A, 5 μm, 50x2.1 mm column (Agilent) and resolved with a gradient from 15% to 50% acetonitrile in water with 0.1% formic acid at 0.5 ml/min flow rate. Positive polarity Q-TOF mass spectra were acquired in m/z range 300-3200. Spectra were deconvoluted using the BioConfirm software (Agilent). The number of phosphate groups was calculated as a multiple of 80.0 Da added to the mass of the unmodified protein.

MALDI-TOF was carried out with a MALDI Rapiflex (Bruker Daltonics, Bremen, Germany). 2 µL protein was mixed with 10 µL matrix consisting of 10mg/mL of sinapinic acid (Sigma Aldrich, St. Louis, MO, cat# 85429), in 50% (v/v) acetonitrile (Thermo Fisher Scientific, Waltham, MA, cat# 26827-0040) and 0.05% trifluoroacetic acid (Sigma Aldrich, cat# T6508), and 2 µL of this mixture was deposited on a MTP 384 ground steel plate. Data were acquired in positive ion mode with linear detection from 10 to 80 kDa at a sample rate of 0.63 GS/s, with 4,000 laser shots, and laser power at 90%.

### Materials and data availability

Plasmids for N protein and mutants used in the biophysical studies will be shared on request. Raw data of all figures and SV-AUC analysis software can be accessed at the Harvard Dataverse (https://doi.org/ [to be assigned]).

## Acknowledgements

This work was supported by the Intramural Research Program of NIBIB (ZIA EB000099-02), NHLBI, NIDDK, and NINDS at the National Institutes of Health (NIH), with additional support (to D.O.M.) from NIGMS (R35-GM118053). The contributions of the NIH authors are considered Works of the United States Government. The findings and conclusions presented in this paper are those of the authors and do not necessarily reflect the views of the NIH or the U.S. Department of Health and Human Services. We thank the Biophysics Resource in the Center for Structural Biology, Center for Cancer Research, NCI at Frederick for assistance with some of the LC-MS studies.

