## Supplemental Methods for "Intramolecular loops control SARS-CoV-2 nucleocapsid protein self-association and nucleic acid binding dependent on phosphorylation"

*Dimerization with conformational change*

This model describes a protein that exhibits two states, one that can undergo reversible dimerization, and a second one that does not. This maps onto the N-protein with its transient LRS-NTD-loop that is inhibitory for LRS oligomerization, considering the constitutive N-protein dimer (connected through a high-affinity interaction in the CTD) as a unit that dimerizes, and the resulting tetramers as dimers of dimers.

In the dimerization-competent state (in absence of an inhibitory loop LRS-NTD-loop) the monomer-dimer equilibrium is described by the dissociation constant

$$K_{d,12} = \frac{m^2}{d} \quad (6)$$

where $m$ and $d$ are the monomer and dimer concentrations, respectively, and $K_{d,12}$ the equilibrium constant. If we allow the monomer to transiently occupy a state $l$ (with LRS-NTD-loop) unable to dimerize, then the apparent equilibrium dissociation constant $K_{d,app,12}$ follows

$$K_{d,app,12} = \frac{(m+l)^2}{d} \quad (7)$$

After rearranging, this leads to the concentration ratio $l/m$

$$\frac{l}{m} = \sqrt{\frac{K_{d,app,12}}{K_{d,12}}} - 1 \quad (8)$$

, which equals the time-average ratio of the fractional time the monomer existing in the dimerization-incompetent state, $f_l$ , relative to the dimerization-competent state, $f_m$ ; $f_l/f_m = l/m$ . With $1 = f_l + f_m$ it follows that

$$f_l = \frac{\sqrt{K_{dapp,12}} - \sqrt{K_{d,12}}}{\sqrt{K_{dapp,12}}} \quad (9)$$

.

*Analysis of competitive nucleic acid binding*

NA ligands will compete with the pSR-NTD-loop for the binding site on the NTD. Formally, this is equivalent to the classical model of a ligand binding to a single site at a receptor with two conformational states, where one state (pSR-NTD-loop) does not bind the ligand. In this case, the ligand binding displays an apparent binding constant (Vega et al, 2015)

$$K_{NA,app} = K_{NA}/(1 + K_{SR}) \quad (1)$$

where $K_{NA}$ is affinity constant for the NA ligand to the susceptible state, and $K_{SR}$ is the equilibrium constant for the pSR-NTD-loop. From measuring $K_{NA}$ as the NA binding affinity in the non-phosphorylated state (without pSR-NTD-loop), and $K_{NA,app}$ as the binding constant measuring in the phosphorylated state (capable of forming the pSR-NTD-loop), rearranging Eq. 1 we can determine $K_{SR}$ as

$$K_{SR} = K_{NA}/K_{NA,app} - 1 = K_{d,NA,app}/K_{d,NA} - 1 \quad (2)$$

On the other hand, since $K_{SR}$ describes the equilibrium between the populations of protein in the conformational states,

$K_{SR}$ = [population with pSR-NTD-loop]/[population without pSR-NTD-loop] (3)

we can determine the time-average fraction of the pSR-NTD-loop configuration

$$f_{pSR-NTD-loop} = K_{SR}/(1 + K_{SR}) \quad (4)$$

Furthermore, the free energy of binding for the pSR-NTD-loop is

$$\Delta G_{SR} = -RTln(K_{SR}) \quad (5)$$

where $R$ is the gas constant and $T$ the absolute temperature.
