## Supplemental Figure for "Intramolecular loops control SARS-CoV-2 nucleocapsid protein self-association and nucleic acid binding dependent on phosphorylation"

### Supplemental Figure S1

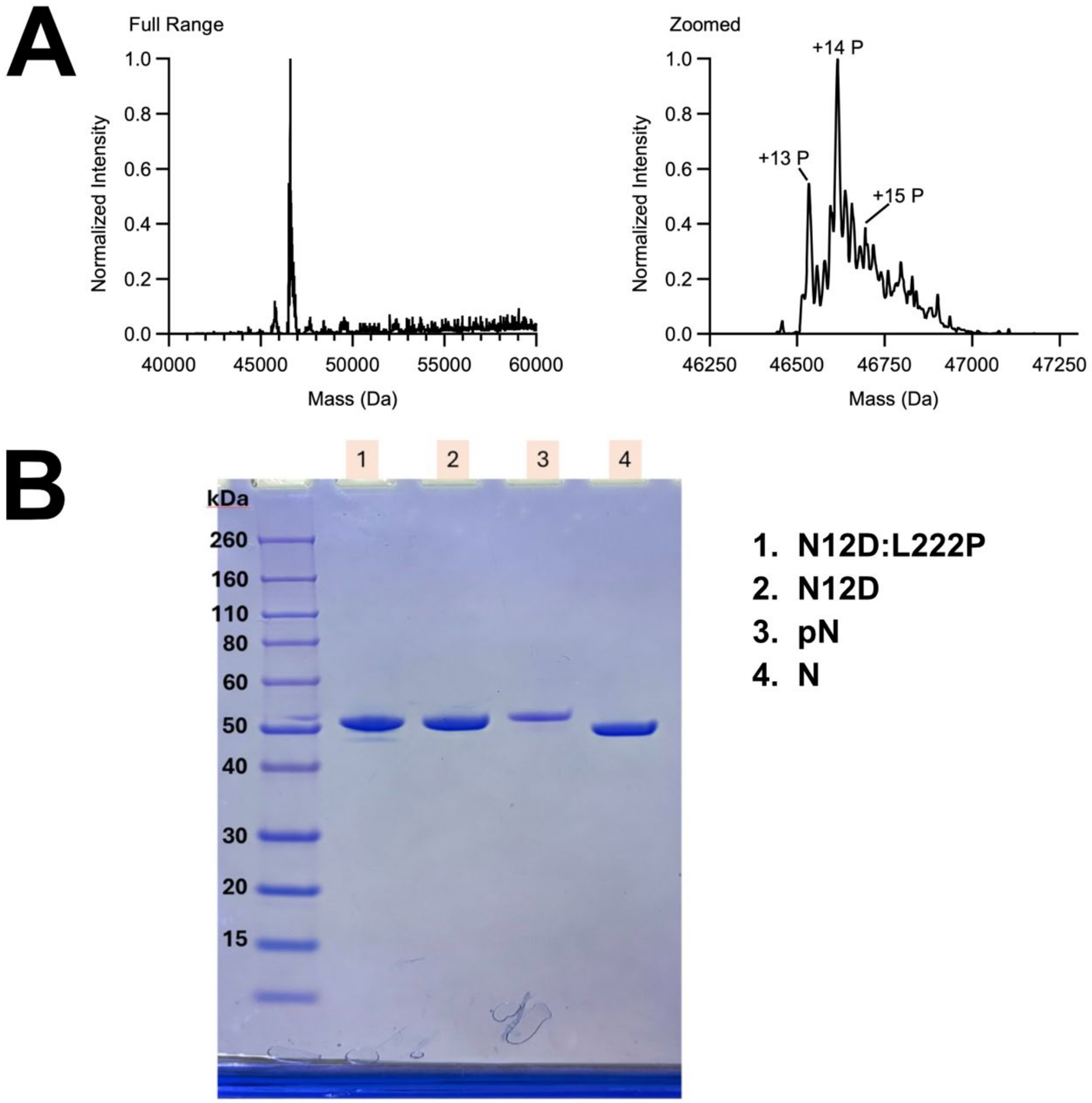


**Supplemental Figure S1: Characterization of enzymatically phosphorylated N**. (**A**) Deconvoluted mass spectra from the LC/MS analysis of N. Molecular weights of 46535, 46616, and 46696 Da correspond to N+13, 14 and 15 phosphate groups, respectively. (**B**) Purity and comparison of mobility in SDS-PAGE of unmodified N, phosphorylated N, phosphomimetic $N_{12D}$ , and phosphomimetic $N_{12D}$:L222P.

### Supplemental Figure S2

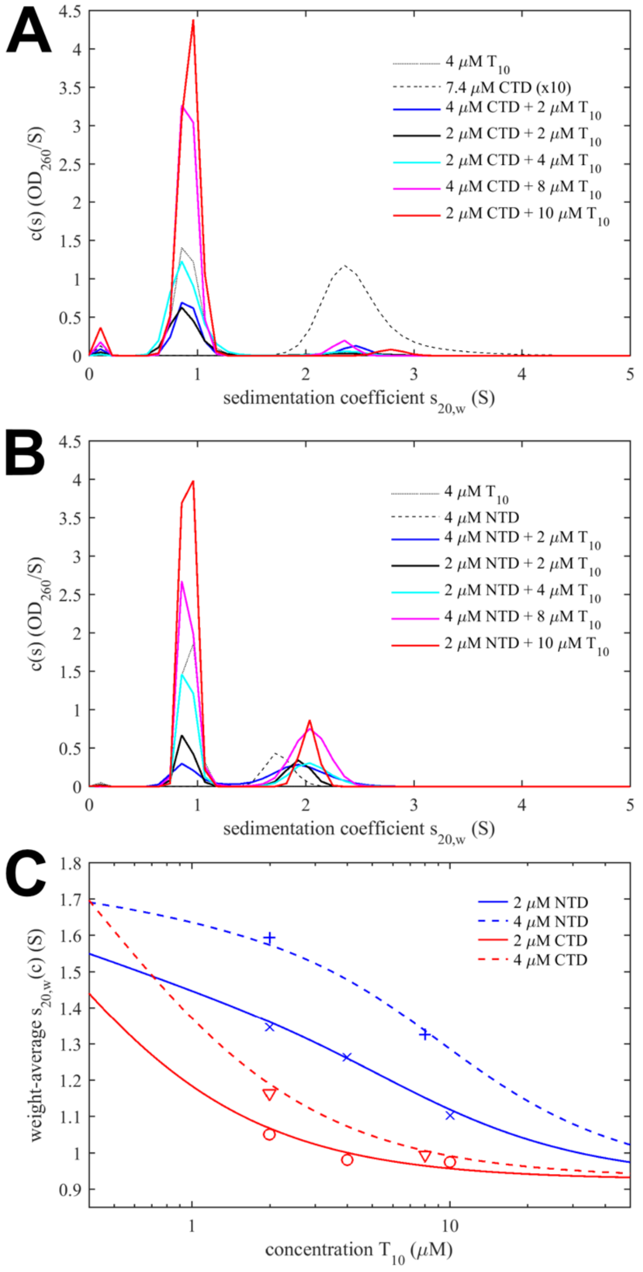


**Supplemental Figure S2: Oligonucleotide $T_{10}$ binds strongly to the NTD and little to the CTD**. (**A**) Concentration-dependent sedimentation coefficient distributions *c*(*s*) for mixtures of CTD ($N_{247\text{-}364}$) and oligonucleotide $T_{10}$ in 20 mM HEPES, 150 mM NaCl, pH 7.5, based on SV-AUC data acquired at 260 nm. (**B**) *c*(*s*) distributions of NTD ($N_{48\text{-}173}$) with $T_{10}$, analogous to (**A**). (**C**) Isotherms of signal-weighted average sedimentation coefficients (symbols) and global fits of a single-site binding model (lines) resulting in a best-fit $K_d$ of 0.69 (> 0.31) mM for the CTD, and 2.9 (2.6 – 4.6) µM for the NTD.

### Supplemental Figure 3

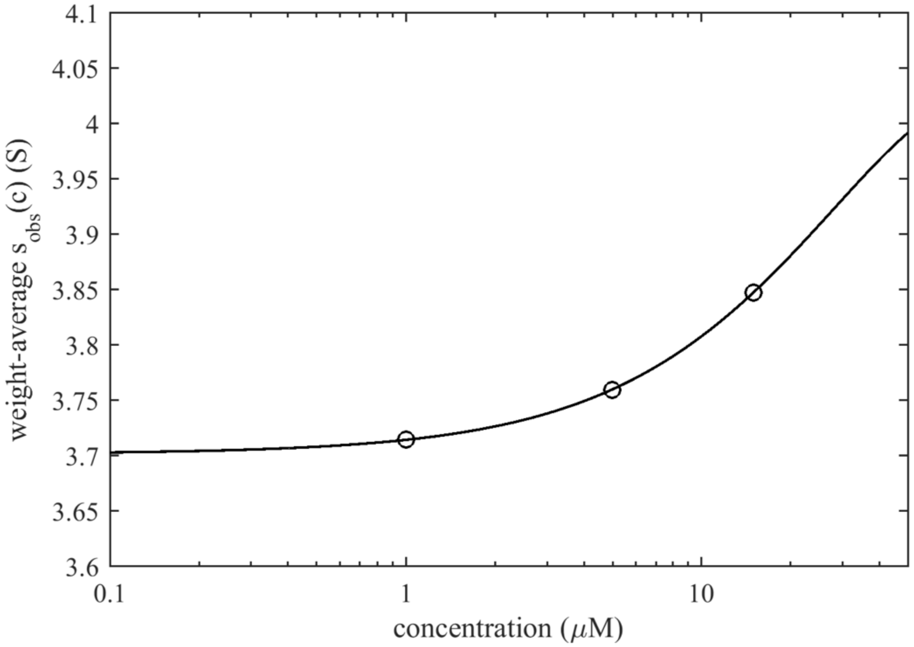


**Supplemental Figure 3: Higher self-association is pH-dependent.** Isotherms of weight-average *s*-values for phosphomimetic $N_{12D}$ in 50 mM sodium phosphate, 150 mM NaCl, pH 6.5 (circles) and best-fit isotherm (line) with a $K_{d,2\text{-}4}$ value of 0.26 mM.

**Supplemental Figure S4**

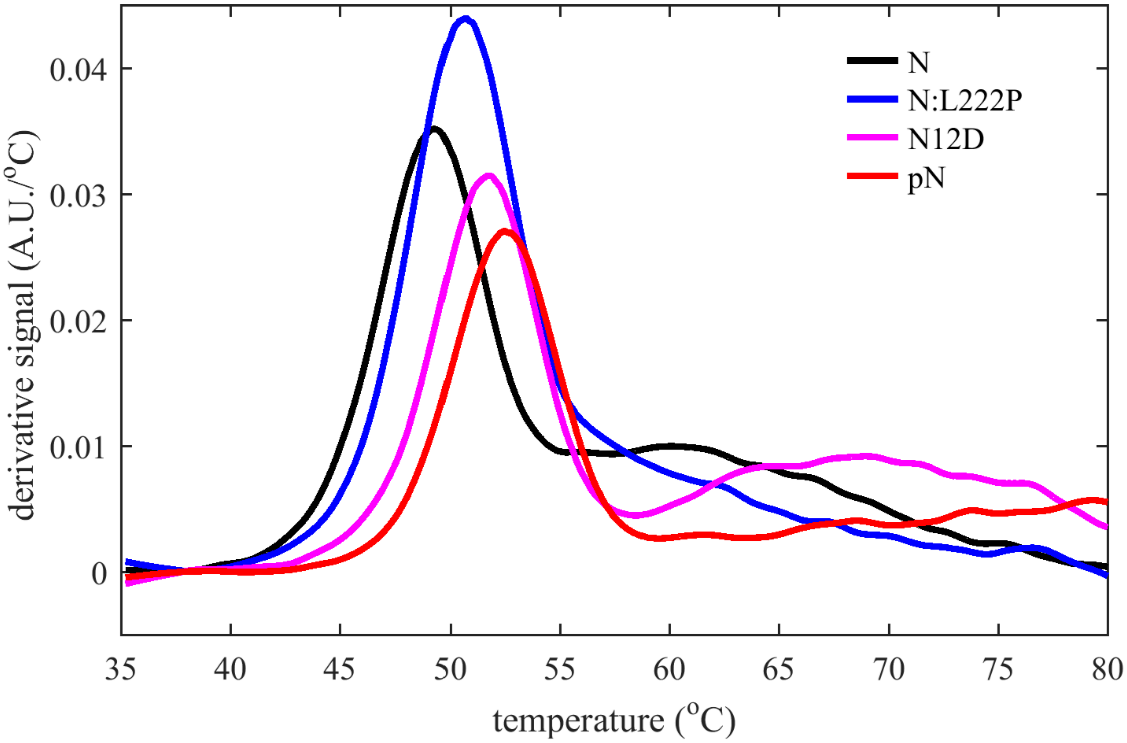


**Supplemental Figure S4: Temperature-dependence change of the intrinsic fluorescence ratio 350 nm and 330 nm depends on intramolecular loops to the NTD.** The main peak reflects solvent exposure of tryptophane and tyrosine residues in the NTD upon melting. (The CTD has a higher melting point of ≈55 °C, and smaller intrinsic fluorescence change due to greater initial surface exposure of its aromatic amino acids; the broad higher transition observed for FL-N coincides with protein aggregation or condensation [18].) Temperature scans for constructs with different phosphorylation state (unphosphorylated N, phosphomimetic $N_{12D}$, and hyperphosphorylated pN) exhibit increasing inflection temperature ($T_i$) with increasing charge in the SR-rich region, reflecting increasing stability of the pSR-NTD-loop. For comparison, the N:L222P mutant abrogates LRS coiled-coils and increases the population of the LRS-NTD-loop to the NTD, thereby also increasing the NTD stability. Scans were acquired with 3 µM protein in 20 mM HEPES, 75 mM KCl, pH 7.4.

### Supplemental Figure S5

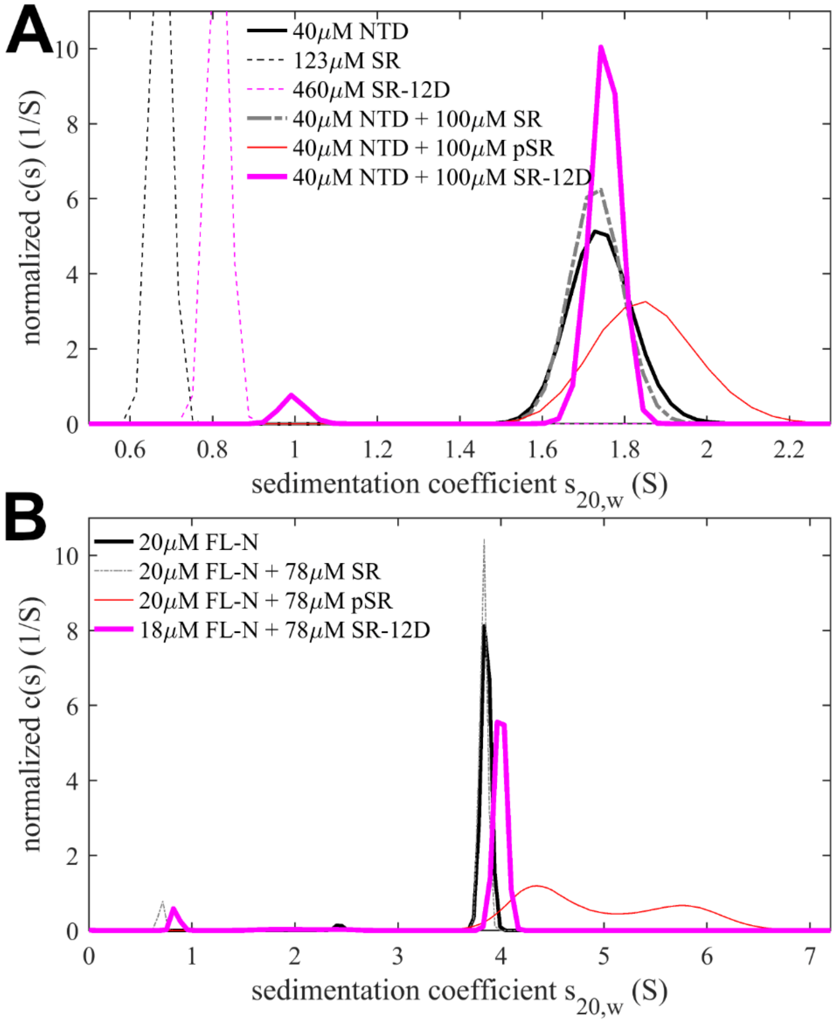


**Supplemental Figure S5: Binding of free SR-12D phosphomimetic peptide to NTD and full-length N in solution.** Analogous to **Figure 2**, SV-AUC experiments were carried out in 20 mM HEPES, 75 mM KCl, pH 7.50, acquiring sedimentation profiles by absorbance at 230 nm, monitoring largely the NTD and FL-N. Sedimentation coefficient distributions were normalized to unit area for comparison of individual components and peptide mixtures with isolated NTD (A) and full-length N (B). The shift of the protein *s*-values in the presence of 12D peptide (magenta) is within error for the NTD, but significant for FL N; in both cases binding of the 12D peptide is much weaker than that of phosphorylated pSR.

### Supplemental Figure S6

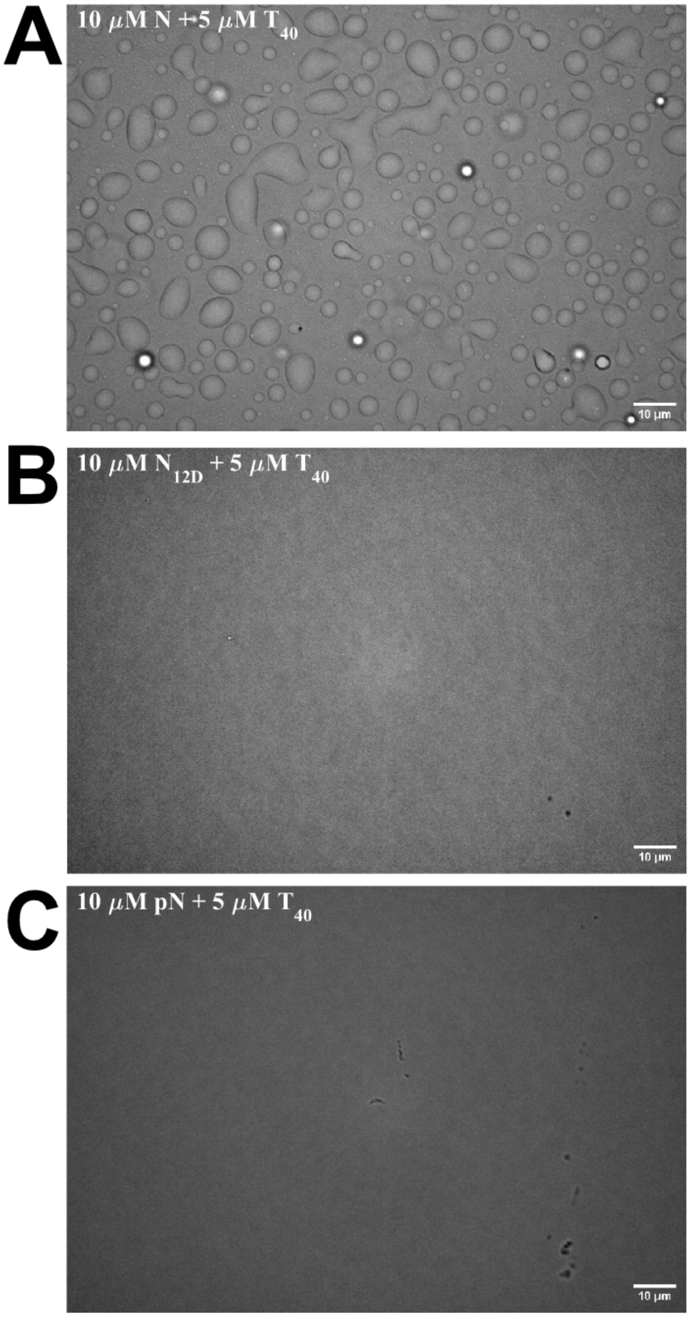


**Supplemental Figure S6: Phosphorylation inhibits macromolecular condensation.** LLPS of unmodified N protein (A), phosphomimetic (B) and phosphorylated N (C) with oligonucleotide $T_{40}$ in 20 mM HEPES, pH 7.5, 10 mM KCl.

**Supplemental Figure S7**

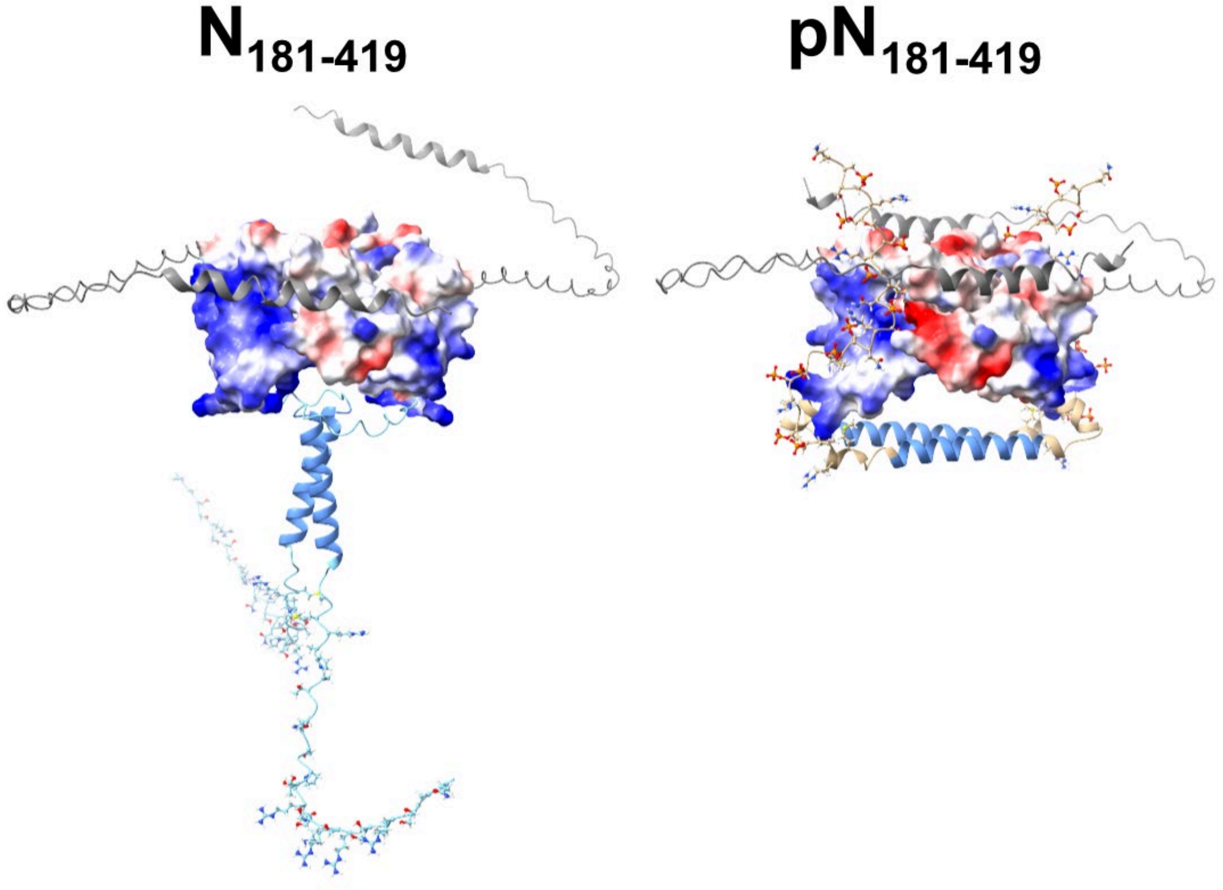


**Supplemental Figure S7: AlphaFold3 prediction of a $N_{181\text{-}419}$ construct lacking the N-arm/NTD suggests possible (non-native) interactions of phosphorylated SR-rich linker region with the CTD dimer.** The CTD dimer is shown with surface rendering, the disordered C-arm as grey cartoon, the LRS helix is shown in blue, and the SR-rich region is shown with atoms. The CTD dimers of both predicted structures are aligned. Without phosphorylation (left) a lack of interactions of the SR-rich region leads to an extended configuration. By contrast, with phosphorylation of the SR-rich region (right), the pSR chains are predicted to make contacts to symmetric groves on the CTD dimer surface.

**Supplemental Figure S8**

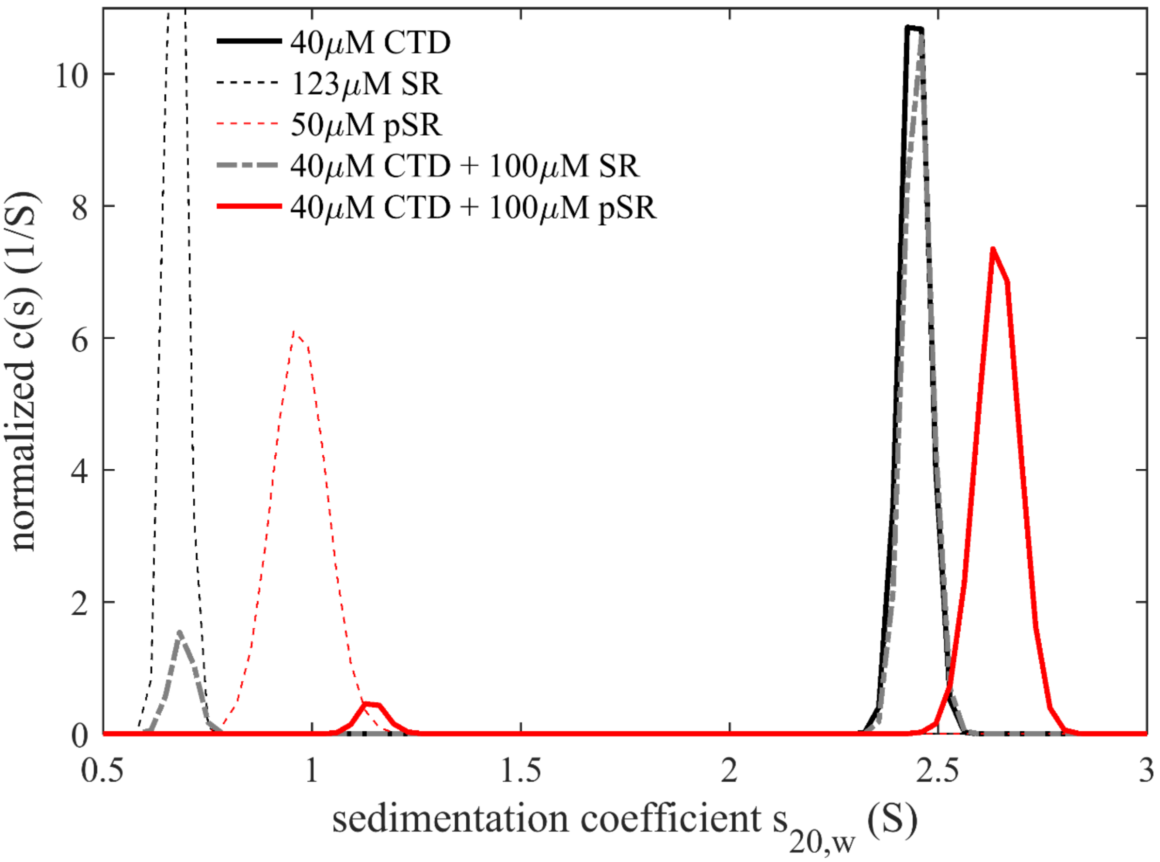


**Supplemental Figure S8: Binding of soluble phosphorylated SR peptide to the isolated CTD in solution.** SV-AUC experiments were carried out analogously to those in **Figure 2**, monitoring the sedimentation of a CTD dimer construct ($N_{247\text{-}364}$, expressed and purified as previously described [22]), phosphorylated or unphosphorylated peptides comprising the SR-rich region ($N_{175\text{-}208}$), and mixtures of peptide with CTD. In the presence of phosphorylated peptide pSR, the CTD peak shifts to a higher *s*-value, indicating complex formation.
